# Frataxin deficiency in proprioceptive neurons is causal to inflammatory and glial responses in dorsal root ganglia

**DOI:** 10.1101/2024.04.16.589410

**Authors:** Pauline Meriau, Laure Weill, Hélène Puccio, Cendra Agulhon

## Abstract

Friedreich ataxia (FA), the most common recessive hereditary ataxia, is an early-onset neurodegenerative disease characterized by pathological changes occurring first in the peripheral dorsal root ganglia (DRG), with loss of the large sensory proprioceptive neurons, leading to ganglionopathy and proprioceptive deficits. FA is caused by a mutation in frataxin gene (*Fxn*), leading to reduced expression of frataxin protein (FXN), an essential ubiquitous mitochondrial protein. Most research has focused on the pathophysiological involvement of proprioceptors. However, in recent years, neuroinflammation is increasingly recognized as an integral and critical contributor in FA pathogenesis. Furthermore, it has also recently been shown a primary reactivity of satellite glial cells (SGCs; glia tightly enwrapping proprioceptor cell bodies), suggesting a role of inflammation and SGC responses in the destruction of proprioceptors in FA patients’ DRGs. It remains unclear to what extent the increase in DRG macrophage response and/or SGC reactivity may contribute to FA phenotype. Therefore, it is important to fully study and understand the mechanism of proprioceptor-macrophages-SGC interactions and their regulations. Exploring relationship between these three cell types has profound implications for breaking through the limitation of treatment of FA. Here we asked whether FXN deficiency selectively in DRG proprioceptive neurons is sufficient to cause inflammatory and glial responses found in patients’ DRG. We used RNA profiling, bioinformatics signaling network and pathway analysis, combined with immunohistochemistry and behavioral experiments to reveal some genes, signaling pathways in macrophages and SGCs that may represent potential biomarkers of the disease. Our study revealed that proprioceptor FXN deficiency causes major changes in inflammatory macrophage and SGC gene transcription as well as macrophage and SGC number, highlighting molecular and cellular pathways that were sequentially altered, thus representing temporal signatures of FA ganglionopathy progression.

## Introduction

Friedreich’s Ataxia (FA) is the most frequent recessive ataxia for which there is no treatment to cure it to date^1^. It is caused by mutations in frataxin gene (*Fxn*), encoding an essential mitochondrial FXN protein, which is involved in iron-sulfur clusters, essential protein cofactors implicated in a large number of cellular functions^2,3^. The most common mutation is an expansion of GAA triplets in the first intron of *Fxn*, resulting in reduced FXN protein expression^4–6^. Moreover, the disease onset is related to FXN expression level, meaning that the higher the number of repetitions is, the earlier the disease onset occurs^5^. This progressive neurodegenerative disease is characterized by pathological changes, occurring first in the peripheral dorsal root ganglia (DRG) with loss of the large sensory proprioceptive neurons^7^. Then, the neurological symptoms are a consequence of degeneration of proprioceptive neurons and their axons in the spinocerebellar tracts as well as neurons in the dendate nuclei and Purkinje cells of the cerebellum^7–9^. *Fxn* gene is ubiquitously expressed, and reduced levels of FXN protein also affect non-neuronal cells and tissues. In particular, most FA patients develop a cardiomyopathy, which is the leading cause of death^5,10,11^.

In DRGs, cell bodies of proprioceptors are tightly enwrapped by satellite glial cells (SGCs)^12^. SGCs carry neurotransmitter receptors and transporters and regulate the neuronal microenvironment homeostasis. Sensory neurons and their surrounding SGCs thus form a distinct morphological and functional unit^13^. Since FA has long been thought to be a strictly neuronal pathology, most research in DRGs has focused on the pathophysiological involvement of proprioceptors. However, recent evidence has shown a primary reactivity of SGCs and suggested a role of inflammation in the destruction of proprioceptors in FA patients’ DRGs. For instance, it has been shown an hypertrophy and proliferation of SGCs, increased level of SGC specific connexin 43 (Cx43)-gap junctions, altered expression of both SGC glutamate transporter GLAST or potassium channel Kir4.1, and increased monocytes entering SGC layers^9^.

Despite the emerging roles of SGCs and macrophages in the pathogenesis of proprioceptors, interactions between these three different cell types are not well understood. Because FXN is a ubiquitous protein, all cell types, including SGC and immune cells, may contribute to proprioceptor dysfunction and degeneration. To start disentangling the respective contribution of each cell type in FA ganglionopathy, here we investigated whether selective FXN depletion in proprioceptive neurons is sufficient to cause inflammatory and SGC responses reminiscent of patients’ phenotype, which both may in return contribute to FA ganglionopathy etiology. We used a conditional mouse model with FXN deficiency specifically in parvalbumin (PV)-expressing neurons, which in DRGs are the proprioceptors. This Pv-cKO-FXN mouse model, recapitulates both sensory ataxia and ganglionopathy associated with FA symptomatic stages^14^. It is therefore a powerful tool to specifically study the impact of proprioceptor FXN depletion in immune and glial cells, while avoiding consequences of generalized FXN deficiency in all other cell types of the peripheral nervous system as well as other body organs. We performed a transcriptomic analysis of DRG from Pv-cKO-FXN model compared to WT mice nCounter RNA profiling combined with bioinformatics network and pathway analysis, as well as immunohistochemical analyses to examine structural proprioceptor association with their enveloping SGCs and neighboring macrophages.

## Materials and methods

### Animals

Littermate wild type (WT) and Pv-cKO-FXN mice were used. Pv-cKO-FXN mouse model was previously described and characterized^14^. Mice were grouped housed (3-5 mice/cage), maintained in a temperature- and humidity-controlled animal facility with a 12-hr light-dark cycle with free access to water and standard rodent chow. We used both males and females, and ensured that control and experimental mice came from the same litters and from different parents. After 7.5 weeks of age when Pv-cKO-FXN started to show obvious ataxic phenotype, moistened chow pellets were added on the cage bottom. All animal procedures were approved by the Université Paris Cité ethical committee (CEEA40, agreement 32473). Animal care and procedures were carried out according to the guidelines set out in the European Community Council Directives.

### Immunohistochemistry, image acquisition and analysis

Animals were transcardially perfused with 4% paraformaldehyde under ketamine/xylazine (100 mg/kg– 10 mg/kg respectively, i.p.) anesthesia. Lumbar L3, L4 and L5 DRGs (*i.e.* 6 DRGs/mouse) were removed, post-fixed for 2 h in 4% paraformaldehyde, respectively. Then, tissues were cryoprotected overnight at 4°C in 0.02 M phosphate buffer saline (PBS, pH 7.4) containing 20% sucrose, and frozen in optimal cutting temperature compound. 14 μm thick sections were cut using a cryostat (Leica), mounted on Superfrost glass slides and stored at - 80°C. The day of the experiment, sections were washed 3 times for 15 min each in 0.02 M PBS. Sections were incubated overnight in 0.02 M PBS containing 0.3% Triton X100, 0.02% sodium azide and primary antibodies (***Supplementary table 1***) at room temperature in a humid chamber. The following day, sections were washed 3 times for 15 min each in 0.02 M PBS, and incubated for 2 h at room temperature with secondary antibodies (***Supplementary table 1***) diluted in 0.02 M PBS containing 0.3% Triton X100 and 0.02% sodium azide. Then, sections were washed 3 times for 15 min in 0.02 M PBS and mounted between slide and coverslip using Vectashield medium containing DAPI (Vector Laboratories). Negative controls, *i.e*. slices incubated with secondary antibodies only, were used to set criteria (gain, exposure time) for image acquisition in each experiment. Image acquisition was performed with an Axio Observer Z1 epifluorescence Zeiss microscope, an ORCA Flash 2.8-million-pixel camera, and a PlanNeoFluar 20x/0.5NA objective. Images were extracted using the ZEN 2011 blue edition software (Zeiss). Measurements were performed using ImageJ software (National Institutes of Health, USA). Analyses were based on 6 DRG slices/mouse, with 10-11 mice/genotype/stages. Experiments and analysis were all performed blind of genotype.

To quantify the number of cells and easily identify the cell body to a specific cell type, we used CD68 markers to label macrophages or GLAST to label SGCs, with DAPI to label nuclei. Neurons larger than 40µm in diameter were considered to be proprioceptors. To quantify GLAST expression levels only in SGCs surrounding large proprioceptors we used GLAST staining to delineate and draw regions of interest around SGCs that selectively envelop proprioceptors. On average, we counted 11 proprioceptors per DRG section. For the analysis of nociceptors, we selected 10 neurons smaller than 30µm in diameter on each DRG section.

At the rate of 6 sections per animal, and 10 or 11 animals per genotype for each stage, in total, we counted macrophages and SGCs around more than 660 proprioceptors and 600 nociceptors per genotype (per experimental stage).

### RNA extraction

Animals were anesthetized with isoflurane and sacrificed. L3, L4, L5 DRGs were quickly harvested (average harvest time for the first DRG was 55 sec and 2 min 50 sec for the sixth DRG), frozen and stored at -80°C. Total RNA from 6 DRGs per mouse was extracted. Samples were homogenized in TRIzol reagent (Invitrogen, Life Technologies) with metal beads and mechanically shacked with a TissueLyser apparatus (Qiagen). The obtained lysate was centrifuged 10 min at 17000 g and 4°C. Then, 1/5 volume of cold chloroform was added to the TRIzol extract, vortexed 30 sec and centrifuged for 15 min at 17000 g at 4°C and the aqueous upper phase was kept. Finally, RNAs were precipitated with 2.5 volume of 100% cooled ethanol and 1µL of glycogen, overnight at -20°C. RNAs were then centrifuged 30 min at 17000 g at 4°C, then washed 3 times with 1mL of 70% cooled ethanol and centrifuged 15 min at 17000 g at 4°C between each wash. Pellets were dried and re-suspended in ultrapure water at room temperature. Protein and TRIzol contaminations were evaluated by measuring their absorbance at 280 and 230nm, respectively. Including criteria were the set as follows: A260/280 and A260/230 values greater than 1.6. Then, RNA integrity was tested using a bioanalyzer and kept for further experiments if the RIN values were greater than or equal to 7. In order for the samples to be comparable, we also defined a DV200% (representing the percentage of RNA with a size greater than 200 nucleotides, which is necessary for probe binding) greater than 60% (***Supplementary table 2***). Analyses based on 6 DRG /mouse, with 4 mice/genotype/experimental stages. Experiments and analysis were all performed blind of genotype.

### RNA transcriptomics

Nanostring procedure (NanoString Technologies, Inc., Seattle, WA, USA) was performed (probe set described in link: <u>Mouse glial profiling gene list</u>) using the *Glial profiling mouse panel* to target 770 genes involved in glial cell biology: cell stress and damage response, pathways regulating glia, inflammation and peripheral immune invasion, glial cell homeostasis and activation, neurotransmission. At least two probes, spanning the exons and 100 bp in size were designed for each gene. One was designed against the 5′-terminal sequence and another against the 3′-terminal sequence. Briefly, probes (a reporter probe and a capture probe) were hybridized to 100 ng of RNA the at 65 °C for 18–24 h using a thermal cycler. Samples were inserted into the nCounter Prep Station to remove excess probes, purify, and immobilize the samples on the internal surface of a sample cartridge for 3 h. Finally, the sample cartridge was transferred to the nCounter Digital Analyzer, where color codes were counted and tabulated for each target molecule. The expression number for the base sequence of the probe part was analyzed and graphed using nSolver Analysis Version 4.0 (NanoString) and Rosalind softwares. Functional analysis and interactions network were made using Ingenuity Pathway Analysis software (IPA, Qiagen) on the total list of genes gathering genes modulated for each stage. All data presented here have been normalized to housekeepping gene expression for each animal and as a ratio comparing experimental animals to control animals.

### Behavior

Prior behavioral tests and from 3 weeks of age, all animals were habituated to experimenter handling. ● Hindlimb extension reflex test: mice were suspended by the tail for 5 sec during three consecutive trials separated of 20 sec each. Abnormal hindlimb extension reflex was determined when hindpaws clasped or showed abnormal positions. An error score of “0” was assigned when hindlimb extension was considered abnormal and a score of “1” when the reflex was not impaired. Scores of the tree trials for each animal were then averaged. ● Two-limb hanging test: animals were habituated to the experimental apparatus 24h before the test day. The setup consisted of a 25 cm-long, 3 mm diameter horizontal bar mounted on two vertical bars and positioned 30 cm above the bench surface. Animals were suspended by the forelimbs and two parameters were measured: time to fall, to grasp hindpaws to the horizontal bar and to reach one of the vertical bars. The maximum time allowed to perform the tasks was 30 sec. A score was assigned for the 3 parameters: i) the falling score (falling between 1-5 sec: 1; between 6-10 sec: 2; between 11-20 sec: 3; between 21-30 sec: 4; >30 sec or not falling: 5); ii) the grasping score (30 minus the time to grasp the bars with hindpaws); and iii) the crossing score (30 minus the time to reach one of the vertical bars). The suspension was repeated for three trials, each separated by a 1 min break (mice return to their cage during the break). ● Static bar test: animals were habituated to the experimental apparatus 24h before the test day. The setup consisted of a 60 cm long, 2.4 cm diameter horizontal bar placed 40 cm above the bench surface. A goal box containing nesting materials was placed at one end of the bar. Mice were placed at the opposite end of the goal box and time to fall, distance traveled, time to reach the goal box was measured. A score was assigned for the 3 parameters: i) the falling score (falling between 1-3 sec: 1; between 4-7 sec: 2; between 8-11 sec: 3; between 12-15 sec: 4; >15 sec or not falling: 5); ii) the distance score was counted as “1” per 10 cm crossed; and iii) the crossing score (30 minus the time to reach one the vertical bars). Tasks were repeated for three trials, each separated by a 1 min break (mice returned to their cage during the break).

Mice were videotaped during experiments. Experiments were performed and movie were analyzed blind of genotype.

### Statistical Analyses

All data are presented as mean ± SEM. Statistical analysis was carried out using GraphPad Prism software (La Jolla, USA). Student’s t tests and ANOVA 2 way were used to compare groups, and a value of p < 0.05 was considered as significant for most experiments, excepted for nCounter RNA results where p < 0.1 was set. Indeed, we reasoned that because proprioceptive neurons represent only ∼11.2% of all DRG sensory neurons (see our quantification in result section), impacts of FXN deficiency in such a low percentage of cells may be difficult to capture at the transcriptome level when considering whole DRGs.

## Results

### Impaired sensorimotor behavior is caused by selective FXN loss in proprioceptors

To further characterized and confirm that the conditional Pv-cKO-FXN mouse model recapitulates sensory ataxia associated to FA^14^, several behavioral tests were performed to evaluate reflex and coordination performances. Compared to WT, Pv-cKO-FXN mice developed a rapidly progressive movement disorder characterized by reflex and coordination defects (***Figure 1*; *Supplementary table 3***). First, hindlimb extension reflex was assessed at five experimental (3.5, 5.5, 7.5, 10.5, and 15.5 week-old) stages, and showed the emergence of reflex abnormalities as early 5.5 weeks of age, which became statistically significantly impaired from 7.5 to 15.5 week-old stages in Pv-cKO-FXN animals compared to controls (***Figure 1a-b*; *Supplementary table 3***). Second, motor coordination performances were scored at 5.5 to 15.5 week-old experimental stages. Two-limb hanging test highlighted a progressive general coordination deficit in Pv-cKO-FXN mice, eventually leading to a dramatic loss of coordination by 10.5 weeks old of age, which was characterized by an increased number of falls, increased difficulty in grasping horizontal bar with hind paws and reaching vertical bars (***Figure 1c-f*; *Supplementary table 3***). Such coordination loss was confirmed by a inability for Pv-cKO-FXN mice to cross the static bar by 10.5 weeks of age (***Figure 1g-j*; *Supplementary table 3***).

**Figure 1.**
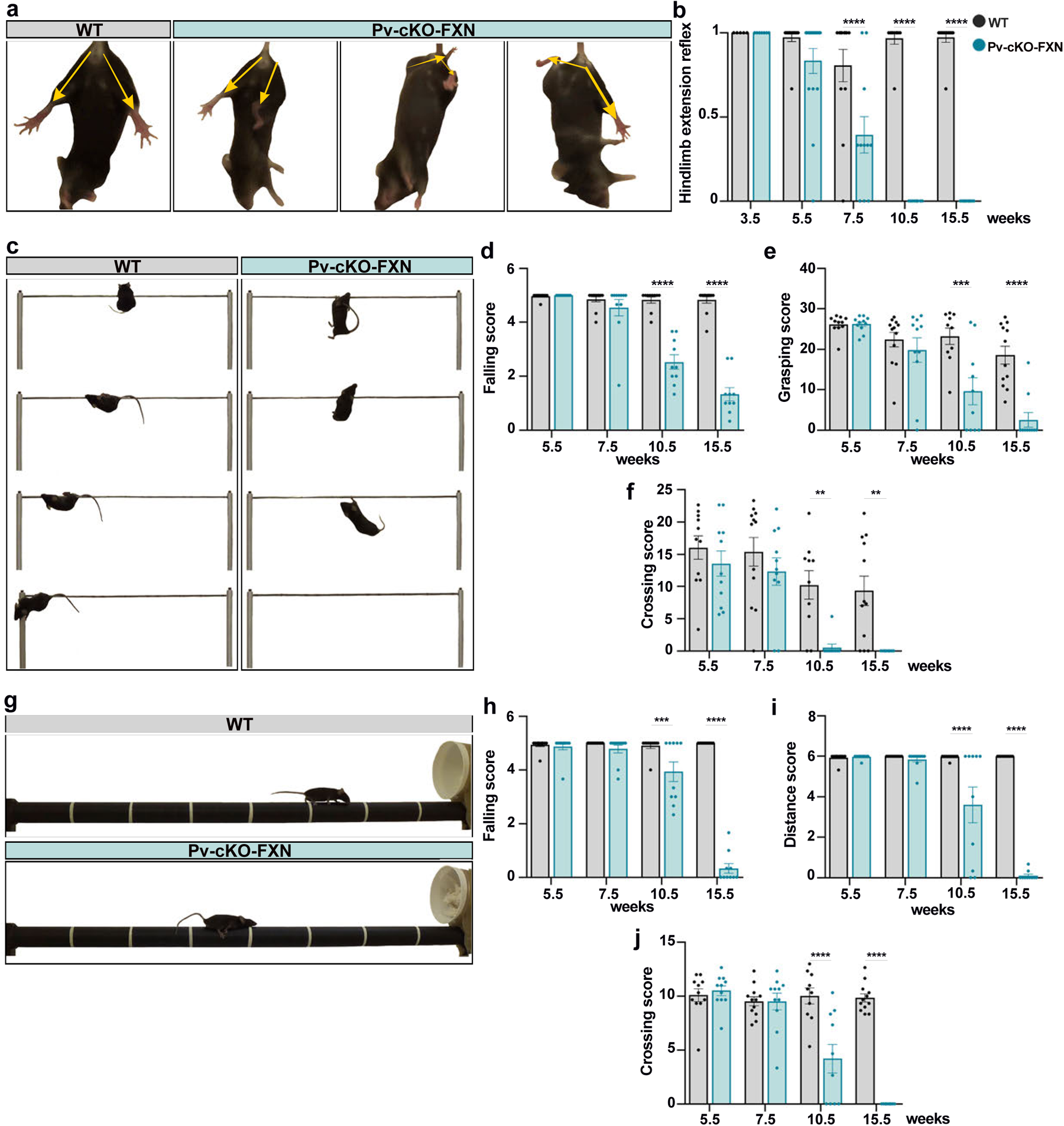
Reflex and sensorimoteurs capacities in Pv-cKO-FXN mice. (**a**) Hindlimb extension reflex in mice (WT, left and Pv-cKO-FXN, right). (**b**) Quantification of abnormal hindlimb extension reflex. (**c**) Two-limb hanging test data. WT mouse (left) grips the horizontal bar and reaches vertical bar. Pv-cKO-FXN mouse (right) shows difficulties to grasping on the horizontal bar and falls. (**d**, **e**, **f**) Quantification of two-limb hanging test data. (**g**) Static bar test data. WT mouse (top) crosses the bar to reach the box. Pv-cKO-FXN mouse (bottom) shows difficulties to cross the bar and falls. (**h**, **i**, **j**) Quantification of static bar test data. Two-way ANOVA compared to WT mice. * p<0.05, ** p<0.01, *** p<0.005, **** p<0.001.

### Selective FXN loss in proprioceptors leads to DRG transcriptional changes and highlights Tspo gene as an early FA biomarker

To determine whether knocking out FXN only in proprioceptors is causal to macrophage and SGC reactivity responses that have been described in FA patients’ DRGs^9^ (*e.g.* enhanced expression of inflammatory and SGC reactivity markers), we performed transcriptomic analysis using nCounter technology. We selected the NanoString *Glial Profiling panel*, which covers 770 genes across 50 pathways involved in glial and immune cell biology throughout both homeostasis and diseases, thus allowing for a comprehensive study of the complex interplay between peripheral immune cells, glial cells and neurons.

Transcriptomic changes were investigated within whole DRGs from Pv-cKO-FXN mice at the five above-mentioned experimental stages, *i.e.* behaviorally asymptomatic 3.5, early symptomatic 5.5, intermediate symptomatic 7.5 as well as the two late symptomatic 10.5 and 15.5 week-old stages, respectively. Among all stages, we identified a total of 76 differentially expressed genes (DEGs) in Pv-cKO-FXN compared to WT mice (***Figure 2a***). Only *Tspo* gene was consistently differentially expressed across the five experimental stages, showing a ∼26-60% up-regulation (***Figures 3-7***). *Tspo* is known to encode an evolutionary conserved protein (TSPO) located in the outer membrane of mitochondria. It is involved in a wide range of mitochondrial functions, including cholesterol transport, steroid hormone and heme biosynthesis, apoptosis and cell proliferation^15^. Notably, TSPO expression level has been found to be increased in different neurodegenerative diseases ^16,17^, including in FA patients’ as well as FA animal model ^18,19^, leading to impairment of mitochondrial antioxidant mechanisms^15^. In clinical practice, TSPO represents a biomarker of neuroinflammation and glial activation^15,20–23^. Thus, *Tspo* up-regulation observed across the five experimental stages in Pv-cKO-FXN mice, not only further validate this mouse model as a powerful FA ganglionopathy model, but also represents a great control for the robustness of our RNA profiling and analysis. Furthermore, *Tspo* is known to be ubiquitously expressed in most DRG cell types, and injured sensory neurons lead to increased *Tspo* expression in macrophages, neutrophils, SGCs, Schwann cell, endothelial cells, and pericytes^24^. Overall TSPO signaling is understood as the result of systemic changes through interactions between various cell types involved in neuroimmune modulation^25^. Therefore our observation that *Tspo* is upregulated in of Pv-cKO-FXN mice, suggest that selective FXN deficiency in proprioceptors may, in itself, trigger complex multicellular responses within DRGs, even before behavioral symptom occurrence (*e.g.* 3.5 weeks of age). This is remarkable given that proprioceptive neurons represent only 11.2% of all lumbar DRG sensory neurons (n = 2,033 proprioceptors among n = 18,194 total sensory neurons; n = 180 L3-L5 DRG slices from WT mice at 7.5, 10.5 or 15.5 week-old stage). Furthermore, because sensory neurons represent themselves 12.5% of all DRG cell types^24^, one can thus estimate that proprioceptors represent ∼1.4% of all DRG cell types, further supporting the idea that FXN deficiency in a very small number of cells, although they are the biggest cells in DRGs, is enough to induce pronounced neuroinflammatory response within DRG.

**Figure 2.**
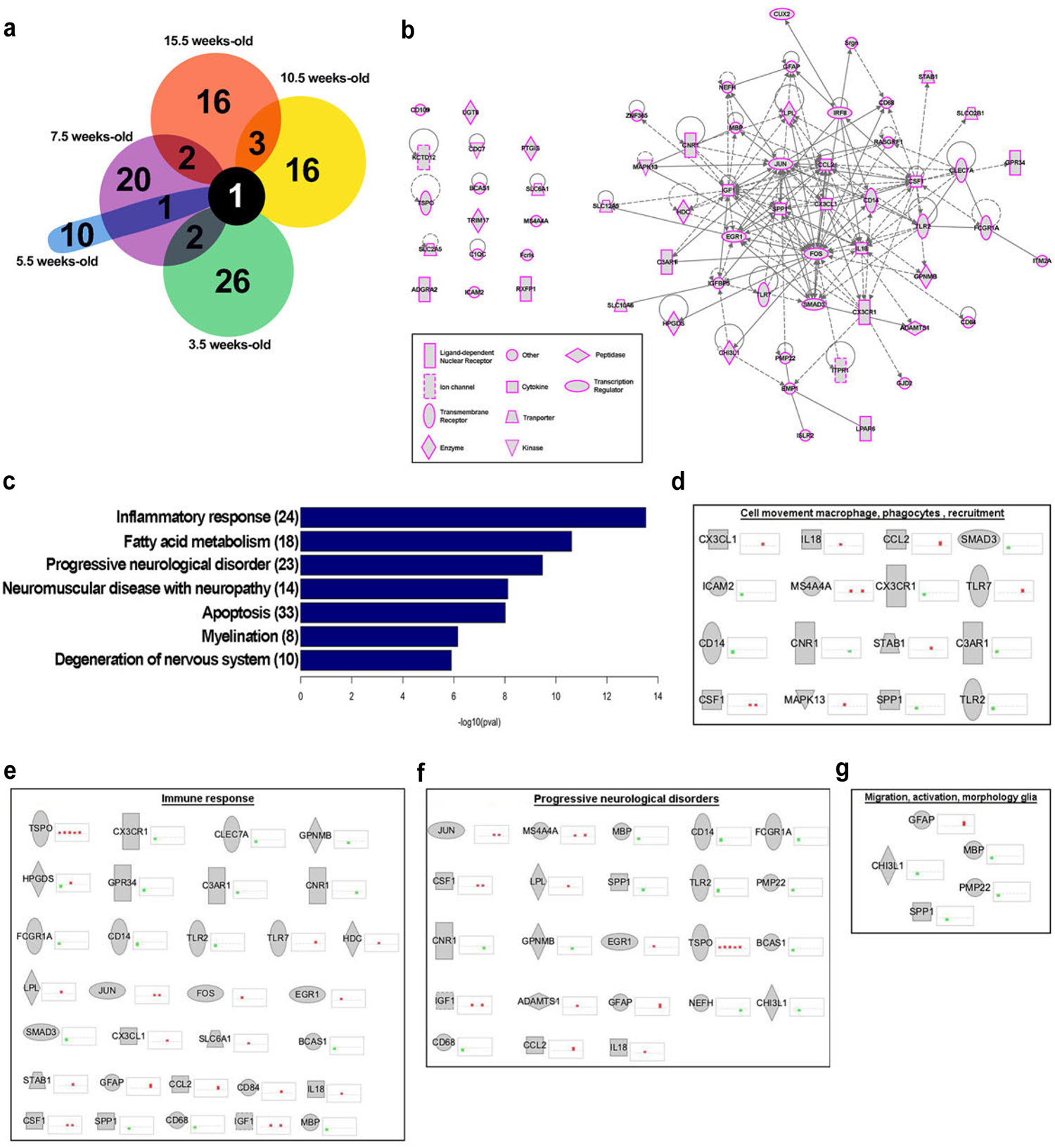
Gene expression analysis of Pv-cKO-FXN mice DRG. (**a**) Venn diagrams comparing signature genes in different stages. (**b**) Networks highlighting the set of differentially expressed genes in the Pv-cKO-FXN model. (**c**) Biological processes analysis (Ingenuity) of enriched genes express Pv-cKO-FXN mice. Functional categories of genes, recruitment of macrophage, phagocytes and cell movement (**d**), immune response (**e**), progressive neurological disorders (**f**) and migration, activation and morphology of glia (**g**). To the right of each gene (displayed in uppercase for clarity) the bars represent the different stages, the color red: upregulation and green: downregulation indicate the extent of differential expression.

**Figure 3.**
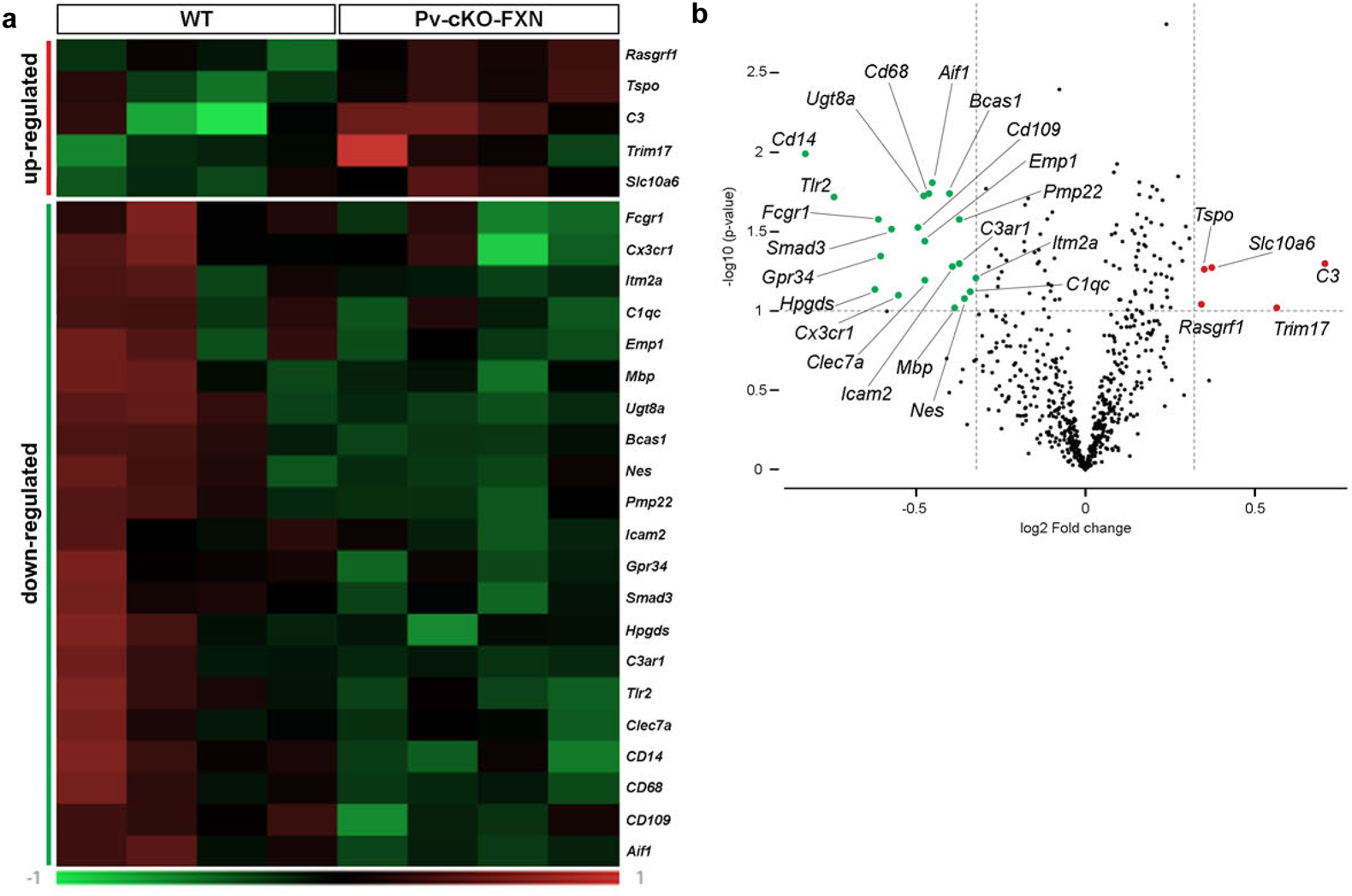
Gene expression analysis of Pv-cKO-FXN mice DRG at 3.5 week-old. (**a**) Heatmap of up- and down-regulated gene (row) for Pv-cKO-FXN compared to WT mice by z-score. (**b**) Volcano plot showing significant differentially expressed genes (fold change -1.25 – 1.25; p-value: 0,1).

Although *Tspo* was upregulated throughout all experimental stages, the vast majority of the 76 DEGs were (up- or down) regulated in a stage specific manner, with only 6% of them that were occasionally shared between 2 stages (***Figure 2a***). This observation is consistent with a progressive, sequential, activation of common signaling pathways. To explore whether some of these DEGs belong to any common molecular network and/or cellular signaling pathway, we performed Ingenuity unbiased functional interaction networks and pathways analysis, taking into account all 76 DEGs, gathering genes modulated in each experimental stage. Importantly, network analysis revealed that 79% of DEGs (*i.e*. 60 genes out of 76; ***Figure 2b***) clustered within a unified molecular network, supporting our hypothesis that these genes, and their gene products may follow common cellular programs in response to proprioceptor FXN knockout. To further determine which shared cellular programs and which cell types may be implicated, we next performed Ingenuity pathway analysis, which revealed a “neuroinflammation signaling pathway” involving 10 (*Tlr2, Tlr7, Cx3cr1, Cx3cl1, Slc6a1*, *Il18, Ccl12*, *Mapk13*, *Jun* and *Fos*) out of the 60 genes as well as two cell types (microglia and astrocytes) (***Supplementary figure 1***). It is worth mentioning that these two cell types were revealed because pathway analysis integrates known data on genomes, chemical molecules, and biochemical systems, including metabolic pathways, drugs, diseases, and gene sequences. However, in the context of our study, macrophages and SGCs should be considered, as they are considered the peripheral counterparts of microglia and astrocytes respectively^26–28^. In conclusion, to our knowledge, our results are the first to support the idea that selective FXN depletion in proprioceptors is causal to DRG TSPO-associated neuroinflammatory responses, involving a dialogue between proprioceptors and, at least, macrophages and SGCs.

To explore whether changes in DEGs were associated with other functions, pathways enrichment analysis was further performed. Our data unraveled 7 significantly enriched functions related to 4 biological processes and 3 disease types. Biological processes included with no surprise inflammatory response (34 genes), but also fatty acid metabolism (18 genes), apoptosis (33 genes), and myelination (8 genes). Disease types included progressive neurological disorder (23 genes), neuromuscular disease with neuropathy (14 genes), and degeneration of nervous system (10 genes) (***Figure 2c***). These 7 functions are all relevant to molecular and cellular phenotypes as well as symptoms observed in FA, supporting further that FXN deficiency in proprioceptors is central in the ganglionopathy pathogenesis.

In an attempt to link FA ganglionopathy with specific genes, we next extracted associations between these 7 functions with i) gene (up or down) regulation across the 5 experimental 3.5, 5.5, 7.5, 10.5 and 15.5 week-old stages, ii) behavioral phenotypes, iii) cell markers, and iv) anatomical cell associations, combined with literature mining. The idea was to identify potential biomarker candidates that have been previously validated for certain cellular or functional phenotypes in order to provide insight into the initiation and progression of FA ganglionopathy. Our main findings are presented in the two following sections.

### FXN loss in proprioceptors caused changes in inflammatory gene transcription, which is accompanied by macrophage and SGC responses

Given the strong evidence for DEGs related to inflammatory processes in DRGs of Pv-cKO-FXN mice (***Figure 2c-e***), we first sought to explore inflammatory gene (and product) functions to have a better understanding of the progression of proprioceptor-induced inflammation. Therefore, we characterized the temporal patterns of DEGs upon FXN knockout in proprioceptors. At 3.5 week-old stage, we observed a total of 26 DEGs in Pv-cKO-FXN mice compared to controls (***Figure 2a***). Among them, 21 genes were down-regulated and 5 were up-regulated (***Figure 3***). 52% of down-regulated genes (11 out of 21 genes, including *Fcgr1*, *C3ar1*, *Gpr34*, *CD14*, *CD68*, *CD109*, *Aif1*, *Tlr2*, *Clec7a*, *Cx3cr1*, *C1qc*) are known to be involved in immune system activation (***Figure 2e***) and to be expressed in DRG macrophages^24^ (Link :<u>DRG</u> <u>profiling</u>). This result suggests a potential impairment in proprioceptor-to-macrophage dialog, leading to delayed immune cell maturation or decreased number of resident macrophages in DRGs of Pv-cKO-FXN mice. In support, concomitant downregulation of *Icam2*, *Itm2a*, *Emp1* genes was also detected (***Figure 3***). These genes are expressed by DRG endothelial cells and SGC^24,29^, and are related to neutrophil extravasation function, transcytosis through the BBB or lymphocyte circulation, respectively^30–33^. As a consequence, expression downregulation of these 3 gene may promote BBB strengthening to prevent infiltration of circulating immune cells into DRGs, and thus preventing secretion of inflammation-associated molecules within DRGs.

Interestingly, at early symptomatic 5.5 week-old stage, expression levels of the 11 (*Fcgr1*, *C3ar1*, *Gpr34*, *CD14*, *CD68*, *CD109*, *Aif1*, *Tlr2*, *Clec7a*, *Cx3cr1*, *C1qc*) gene involved in immune system activation showed comparable levels in Pv-cKO-FXN and WT mice, consistent with the idea of a restored macrophage density (or maturation) in DRGs of Pv-cKO-FXN mice. However, among the 10 DEGs observed at this 5.5 week-old stage (***Figure 4***), three of them (*Itgam*, *Spp1, Chil1*) known to promote immune cell recruitment, were downregulated. *Itgam*, *Spp1* and *Chil1* are expressed by DRG macrophages and/or SGCs^24,29,34,35^ and act as integrin, extracellular matrix remodeling and proinflammatory cytokines^34–38^. Their downregulation thus suggests that immune system and cell invasion might be still attenuated to some extents at this early symptomatic stage in Pv-cKO-FXN *versus* WT mice. By contrast, this same stage is marked by a 58% and 45% increase in transcript expression levels of two immediate early gene transcription factors (*Fos*, *Erg1*) (***Figure 4***). In physio-pathological conditions, both these genes are expressed by DRG macrophages, SGCs, but also endothelial cells, Schwann cells, pericytes, smooth muscles and connective tissues^24,29,39^, and *Egr1* was also reported to be upregulated in DRG throughout disease progression in the ubiquitous FA mouse model above cited^40^. Notably, at this 5.5 week-old stage we also observed the highest (60%) *Tspo* upregulation. All together our findings strongly support the idea that this 5.5 week-old stage is a key turning stage in FA ganglionopathy progression, characterized by a TSPO-associated upregulation of *Fos* and *Erg1* transcription factor gene products, likely leading to a multicellular program activation (*e.g.* involving macrophages, SGCs and endothelial cells). This may consequently induce the modulation of other genes, which effects will be visible at later experimental stages, thus contributing to disease progression. This is exemplified by the emergence of reflex deficits at 5.5 week-old, which became more pronounced at 7.5 weeks of age in Pv-cKO-FXN compared to WT mice (***Figure 1a,b***).

**Figure 4.**
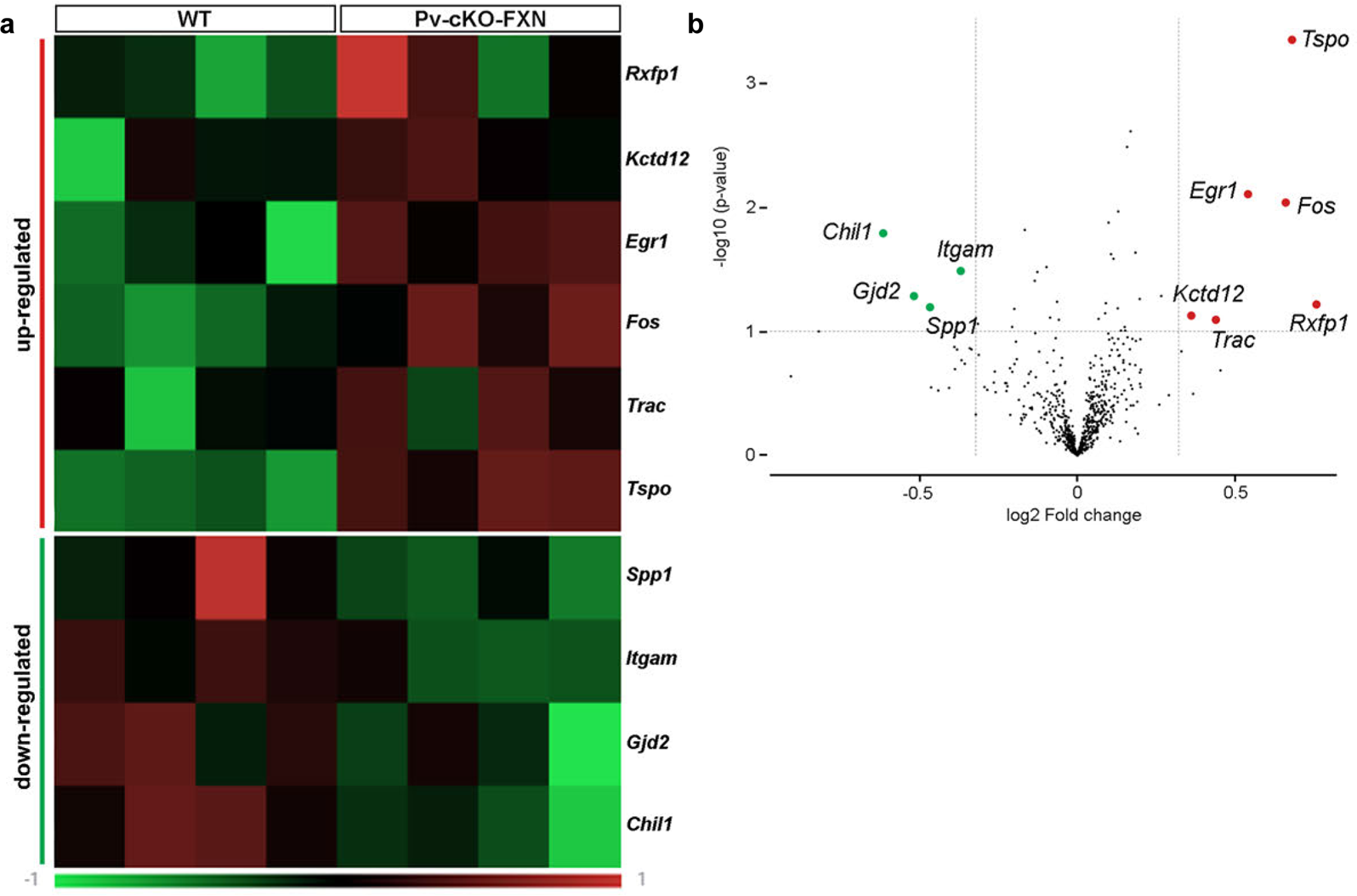
Gene expression analysis of Pv-cKO-FXN mice DRG at 5.5 week-old. (**a**) Heatmap of up- and down-regulated gene (row) for Pv-cKO-FXN compared to WT mice by z-score. (**b**) Volcano plot showing significant differentially expressed genes (fold change -1.25 – 1.25; p-value: 0,1).

Such hypothesis is further supported by the reversal in the ratio of genes that were down-*versus* upregulated at later 7.5 week-old intermediate symptomatic stage. We observed 20 DEGs, among them only 4 were downregulated while the majority (16 genes) were upregulated in Pv-cKO-FXN compared to WT mice (***Figure 5***) Notably, 7 of the upregulated genes (*Ms4a4a, Fcrls, Slco2b1, Ptgis, Hdc, Hpgds, Igf1*) are macrophage markers^24,41–47^ , with *Ptgis* known to promote macrophage M2 polarization and *Ms4a4a*, *Fcrls* and *Igf1* known to be a signature of M2 macrophages^41,42,44,45,48^. Importantly, *Igf1* upregulation was described in other FA animal models^19,49^ (a mouse and a fly models) as well as in human cerebral ataxia^50^. Together these findings suggest an increase in the number of type M2 macrophages in DRGs of Pv-cKO-FXN, which may thus promote inflammation resolution rather than a pro-inflammatory response^42,51^. In agreement we observed upregulation of genes known for their implication in extracellular matrix remodeling (*Adamts1, Srgn, Igfbp5* ; ***Figure 5***) and endothelial cell permeability (*Emp1*), which all may elicit immune cell extravasation and infiltration into DRGs^33,52–55^. The expression of *Adamts1* and *Igfbp5* in DRG endothelial cells and/or pericytes as well as the expression of *Srgn* in endothelial cells and macrophage^24^ further support the idea of an increased influx of macrophages. To determine whether macrophage density was indeed enhanced upon proprioceptor FXN deficiency, we performed immunohistochemistry using the general macrophage CD68 lysosomal marker in order to label the majority of macrophages. At 7.5 week-old intermediate symptomatic stage, total area of CD68 immunostaining signal increased by 55% in whole DRGs of Pv-cKO-FXN *versus* WT mice, likely reflecting an increased number of macrophages, which is supported by trend increases (28%, 26%, 58%) in macrophage density within entire DRGs or specifically around proprioceptors or nociceptors (***Figure 8*; *Supplementary Table 4***). These observations confirm that, at 7.5 week-old intermediate symptomatic stage, FXN deficiency in proprioceptors causes macrophage invasion throughout DRGs, which correlate with significant behavioral symptoms (***Figure 1b***).

**Figure 5.**
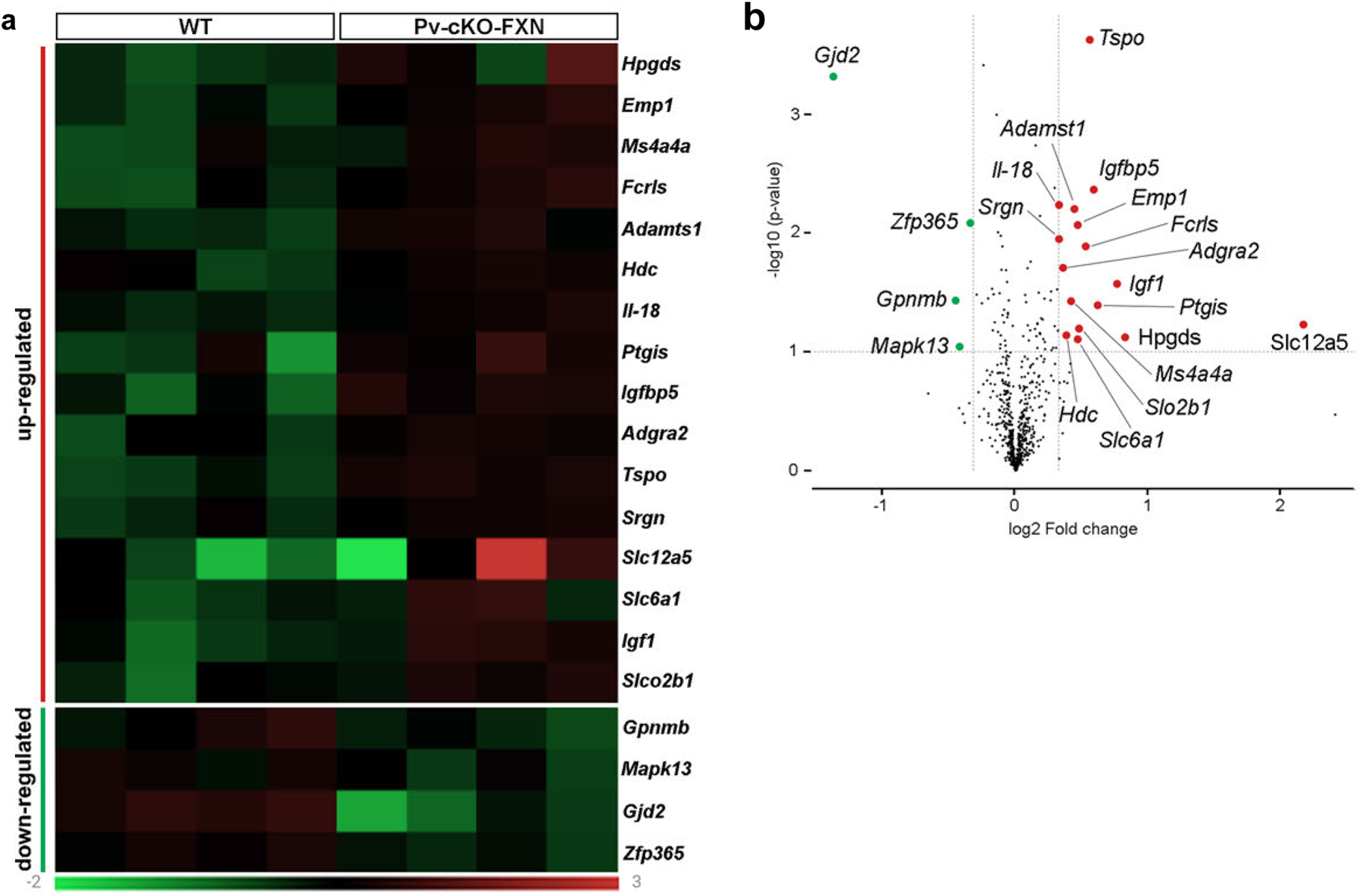
Gene expression analysis of Pv-cKO-FXN mice DRG at 7.5 week-old. (**a**) Heatmap of up- and down-regulated gene (row) for Pv-cKO-FXN compared to WT mice by z-score. (**b**) Volcano plot showing significant differentially expressed genes (fold change -1.25 – 1.25; p-value: 0,1).

To further investigate the progression of inflammatory response and its relation to the severity of reflex and coordination impairments, transcriptomic analysis was performed at 10.5 week-old late symptomatic stage. We observed 16 DEGs, and as for 7.5 week-old stage. Only 3 were downregulated while most of them (13 genes) were upregulated in Pv-cKO-FXN compared to WT mice (***Figure 6***). A majority of upregulated genes are known to have inflammatory functions including *Cx3cl1, Cd84, Lpl, Irf8, Siglech, Lrrc25, Mafb, Stab1,* and *Sphk1* ^24,56–62^. However, here they segregated into two categories: genes related to “alternatively” activated M2 macrophages (as found in previous experimental stages) and genes related to “classically” activated M1 macrophages. *Stab1* and *Mafb* encode scavenger receptor and transcription factor, respectively. They are expressed by M2 macrophages and both promote phagocytosis of apoptotic cells, thus maintaining tissue homeostasis^24,29,61,63–65^. *Irf8, Lpl, Cd84* genes are regarded as important regulators of macrophage differentiation, function as well as polarization of inflammatory M1 macrophage^56,66–69^ , and are expressed in DRG macrophages^24^. Furthermore, *Sphk1* gene transcript was also upregulated but, as opposed to *Stab1, Mafb Irf8, Lpl, and Cd84*, it is expressed in DRG SGCs and not in macrophages^24^ (***Figure 6***). However, at the functional level it also contributes to stimulate proinflammatory responses and M1 macrophage activation^42,70^. Very importantly, we observed an upregulation of two neuronal genes (*Cx3cl1* and *Csf1*)*. Cx3cl1* is known to encode a potent chemokine CX3CL1 (also called Fractalkine). *Csf1* encodes a cytokine (colony-stimulating factor 1; CSF1) and SGC activation can induce *Csf1* upregulation in sensory neurons under pathological conditions^71,72^. In DRGs, both neuronal CX3CL1 and CSF1 acts as chemoattractant for macrophage recruitment, thus representing critical inflammatory mediators involved in neuron-to-macrophage signaling^73–78^. Finally, this 10.5 week-old (as well as the following 15.5 week-old) stage are characterized by an upregulation of *Jun* transcripts (***Figure 6***). *Jun* encodes an immediate early gene transcription factor and represents a signature of cell activation and intense gene transcription. Given that *Jun* is expressed in all DRG cells^24^, such *Jun* upregulation may reflect a general multicellular activation leading to the expression of new genes related to FA pathogenesis (*e.g.* SGC activation, proprioceptor CX3CL1 and CSF1 production and release, macrophage M2 migration and M1 polarization as well as other cellular functions participating to ganglionopathy progression). Taking together, these findings indicate that neuron-SGC-macrophage interactions have changed significantly at 10.5 week-old symptomatic stage compared to earlier 7.5 week-old stage, letting emerge the following possible scenario. At 10.5 week-old stage, FXN deficient proprioceptors have become quite dysfunctional as evidenced by cellular abnormalities^14^ (*e.g.* vacuolization, mitophagy, endoplasmic reticulum dilatation) and drastic deficits in reflex and coordination (***Figure 1***). Surrounding SGCs have sensed proprioceptor activity dysfunction through their numerous neuroactive agent receptors and transporters^13^, have influenced in turn neighboring proprioceptor production and secretion of CX3CL1 and CSF1, and have themselves produce mediators stimulating proinflammatory responses. Diffusion of these mediators through DRGs has subsequently induced i) macrophage recruitment around proprioceptors, and ii) polarization of some immune cells into M1 macrophages to undergo transitions toward proinflammatory functions.

**Figure 6.**
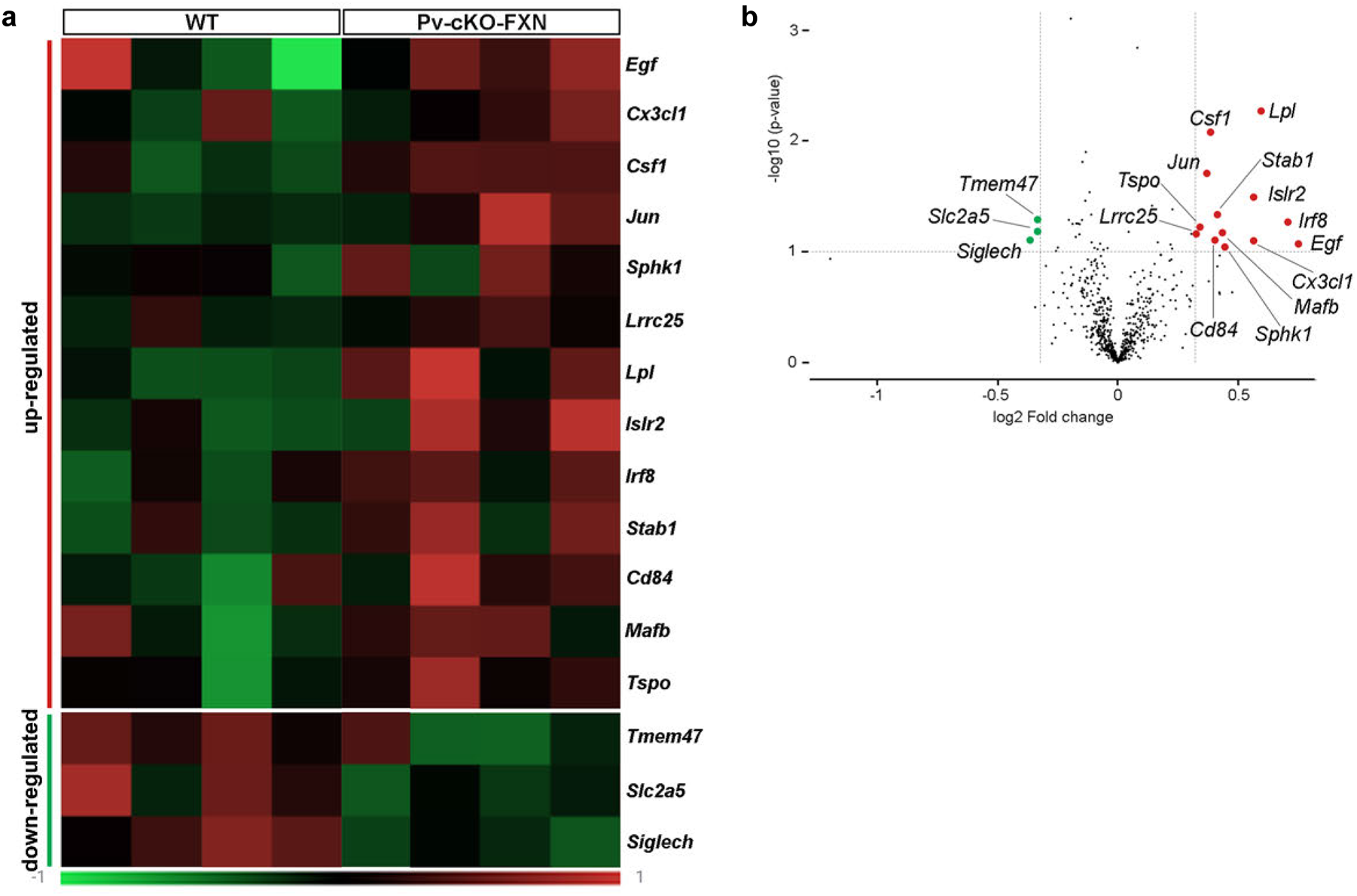
Gene expression analysis of Pv-cKO-FXN mice DRG at 10.5 week-old. (**a**) Heatmap of up- and down-regulated gene (row) for Pv-cKO-FXN compared to WT mice by z-score. (**b**) Volcano plot showing significant differentially expressed genes (fold change -1.25 – 1.25; p-value: 0,1).

**Figure 7.**
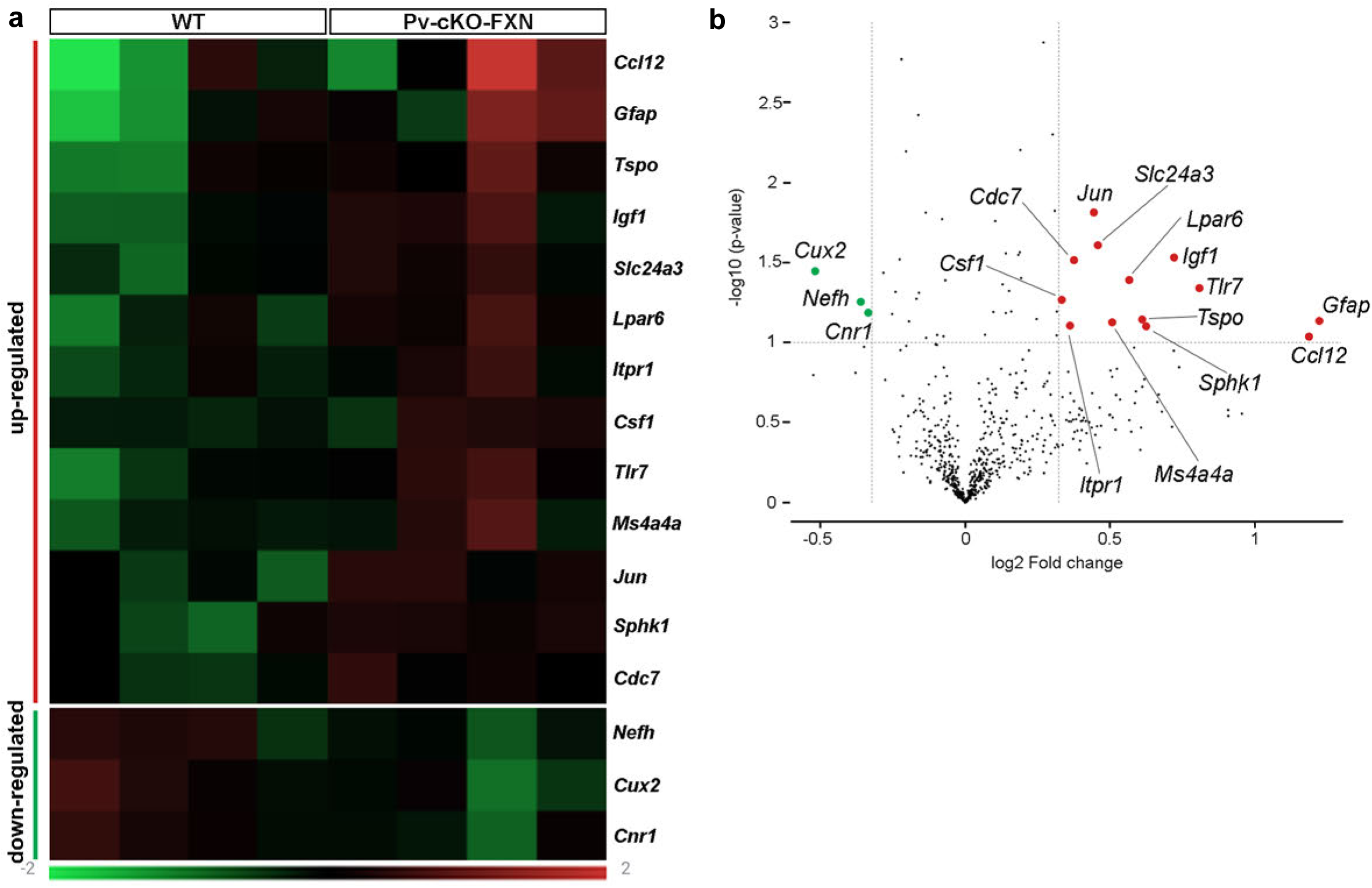
Gene expression analysis of Pv-cKO-FXN mice DRG at 15.5 week-old. (**a**) Heatmap of up- and down-regulated gene (row) for Pv-cKO-FXN compared to WT mice by z-score. (**b**) Volcano plot showing significant differentially expressed genes (fold change -1.25 – 1.25; p-value: 0,1).

Although this scenario remains to be further tested, we assessed whether macrophages were preferentially clustered around proprioceptors. As described above we used immunohistochemistry to detect and quantify CD68 expressing macrophages. Our data showed that the 55% increase in CD68 expression within whole DRGs of Pv-cKO-FXN mice at 7.5 week-old stage was not observed at late 10.5 and 15.5 week-old stages (***Figure 8*; *Supplementary table 4***). However, when quantifying macrophages around the SGCs that enwrap proprioceptors (but not other sensory neurons, *e.g.* nociceptors), we observed an increase in 53% of macrophages at late (10.5 week-old) in Pv-cKO-FXN mice compared to controls (***Figure 8c***). This was not the case for nociceptors (***Figure 8d***), confirming that, at 10.5 week-old stage, macrophages do migrate and accumulate around FXN deficient proprioceptors and their respective surrounding SGCs when major reflex and coordination deficits are observed^14^ (***Figure 1***). This is in agreement with findings from patient’s DRG showing accumulation of macrophages around unhealthy sensory neurons, likely proprioceptors^9^. Then, to determine whether, at this 10.5 week-old stage, macrophages mediate phagocytosis of proprioceptors, we quantified the density of all DRG sensory neurons *versus* the densities of proprioceptors or nociceptors using NeuN staining. These measurements revealed no obvious decrease in neuron number (n = 2 033 proprioceptors and n = 1 800 nociceptors among n = 18 194 total sensory neurons; n = 180 L3-L5 DRG slices, n = 30 WT mice, 10 mice per experimental 7.5, 10.5 or 15.5 week-old stage) (***Supplementary table 4***). Thus, we next sought to test whether macrophages target and phagocyte SGCs that envelop proprioceptors in ganglionopathy progression. To address this question we investigated whether the two SGC specific GLAST and Cx43 protein markers were differentially expressed in Pv-cKO-FXN *versus* WT mice at 10.5 week-old in comparison to 7.5 and 15.5 week-old stages. We detected no changes (albeit a trend decrease) in GLAST expression level within entire DRGs (***Figure 9b*; *Supplementary table 5***). However, we observed a significant 19% decrease in Cx43 expression level within whole DRGs of Pv-cKO-FXN mice at 10.5 week-old stage (***Figure 10b*; *Supplementary table 4***). Given that we found that proprioceptors represent only ∼11.2% of all lumbar DRG sensory neurons and that SGCs exhibit very small cytoplasm and extremely fine layers, it remains a challenge to detect protein expression changes in the subpopulation of SGCs that selectively envelop proprioceptors. In an effort to address further our question, we quantified the number of SGCs surrounding large proprioceptors. It required using GLAST staining to delimit and draw regions of interest around SGC that enwrap proprioceptors selectively. This approach showed that SGCs enveloping specifically proprioceptors are 15% less numerous per proprioceptor at 10.5 week-old symptomatic stage compared to the 7.5 and 15.5 week-old stages (**Figure 9c**), corroborating the decreased in Cx43 expression level observed at 10.5 week-old stage (***Figure 10b***). This was not found in SGCs surrounding nociceptors, further supporting the idea that this decreased in SGC number was specific of the SGC subpopulation enveloping FXN deficient proprioceptors. Furthermore, our results in WT mice showed that the number of SGCs/proprioceptor, quantified on 14 um thick DRG sections, was of 3.66, 3.58, and 3.14 at 7.5, 10.5 and 15.5 week-old stages, respectively (***Supplementary table 5***). These results suggest that, during young adulthood *(i.e.* between 10.5 and 15.5 week-old stages), there is a physiological diminution of SGCs enwrapping large sensory neurons, which was not the case for small nociceptors neurons (***Figure 9b*; *Supplementary table 5***). Because this reduction in SGCs/proprioceptor appears earlier in Pv-cKO-FXN mice (10.5 instead of 15.5 week-old) compared to WT mice, it may have profound implications in ganglionopathy progression. Together our findings suggest that, at 10.5 week-old stage, when macrophages are recruited around FXN deficient proprioceptors, they may phagocyte some SGCs, possibly contributing to (or exacerbating) neuronal microenvironment homeostasis alteration, proprioceptor dysfunction and degeneration, and ultimately sensorimotor deficits.

**Figure 8.**
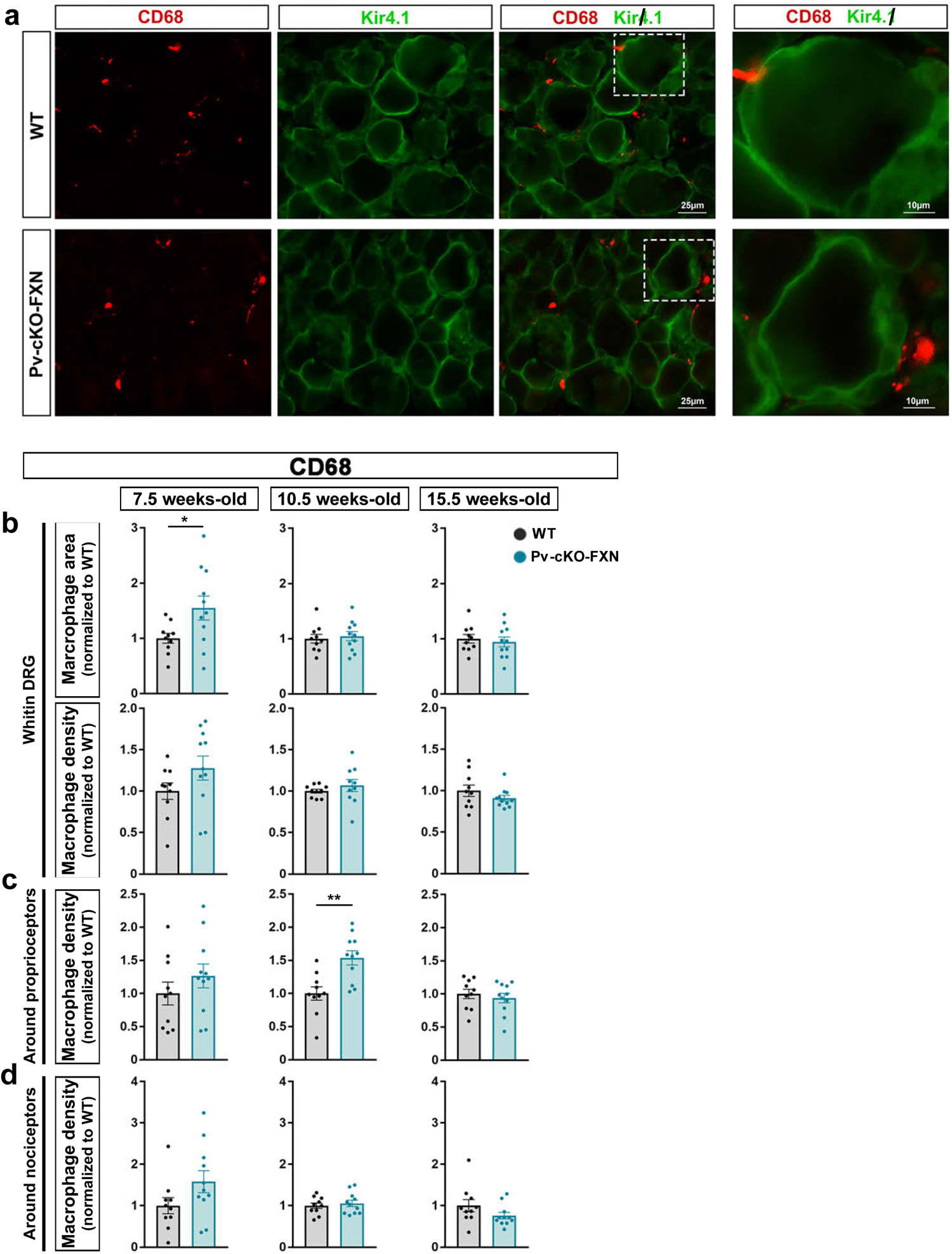
Immunohistochemistry of CD68 and Kir4.1 in DRG of control and Pv-cKO-FXN mice. **(a)** Representative images of immunofluorescence staining of DRG of WT and Pv-cKO-FXN animals at 10.5 weeks of age; with CD68 (red) which labels macrophages in close proximity to SGC marked with Kir4.1 (green). Scale bar: 25µm and 10µm for magnification. (**b**) Quantification area measurements for CD68-immunostaining and the number of CD68+Dapi+ cells across in whole DRG. Quantification of the number of CD68+Dapi+ cells surronding (**c**) large sensory neurons (diameter than or equal to 40µm) and (**d**) small sensory neurons (diameter between 15 and 30µm). All data shown symptomatic stages (7.5, 10.5 and 15.5 week-old); are mean ± SEM. An unpaired parametric T-test has been performed. ***p<0.005; n=10 WT mice; n=11 Pv-cKO-FXN mice.

**Figure 9.**
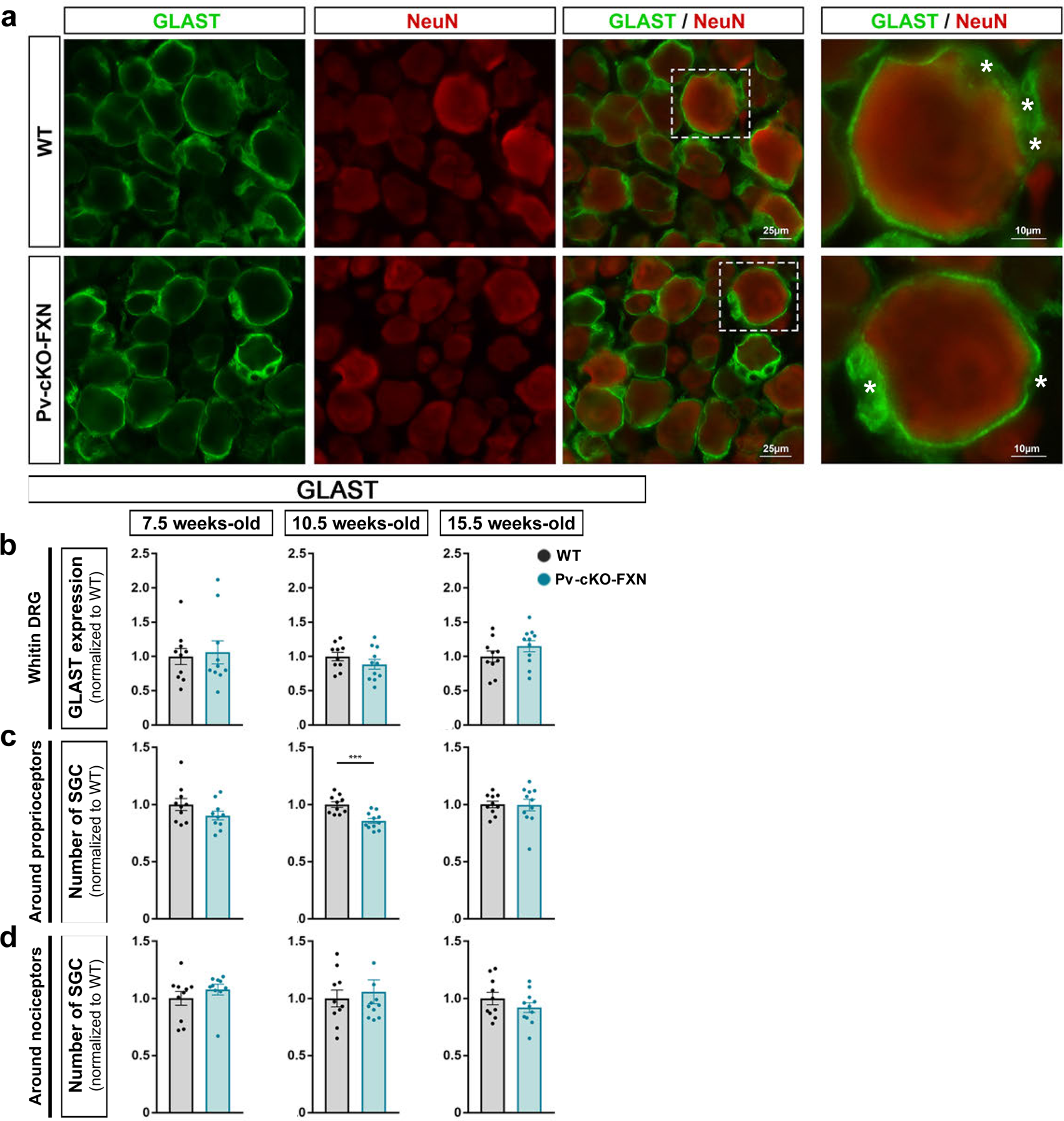
Immunohistochemistry of GLAST and NeuN in DRG of control and Pv-cKO-FXN mice. (**a**) Representative images of immunofluorescence staining of DRG of WT and Pv-cKO-FXN animals at 10.5 weeks of age; with GLAST (green) which labels SGC surrondding neurons marked with NeuN (red). Scale bar 25µm and 10µm for magnification. (**b**) Quantification mean density measurements for GLAST-immunostaining across the entire DRG. Quantification of the number of GLAST+ cells surronding (**c**) large sensory neurons (diameter than or equal to 40µm) and (**d**) small sensory neurons (diameter between 15 and 30µm). All data shown symptomatic stages (7.5, 10.5 and 15.5 week-old) are mean ± SEM. An unpaired parametric T-test has been performed. ***p<0.005; n=10 WT mice; n=11 Pv-cKO-FXN mice.

**Figure 10.**
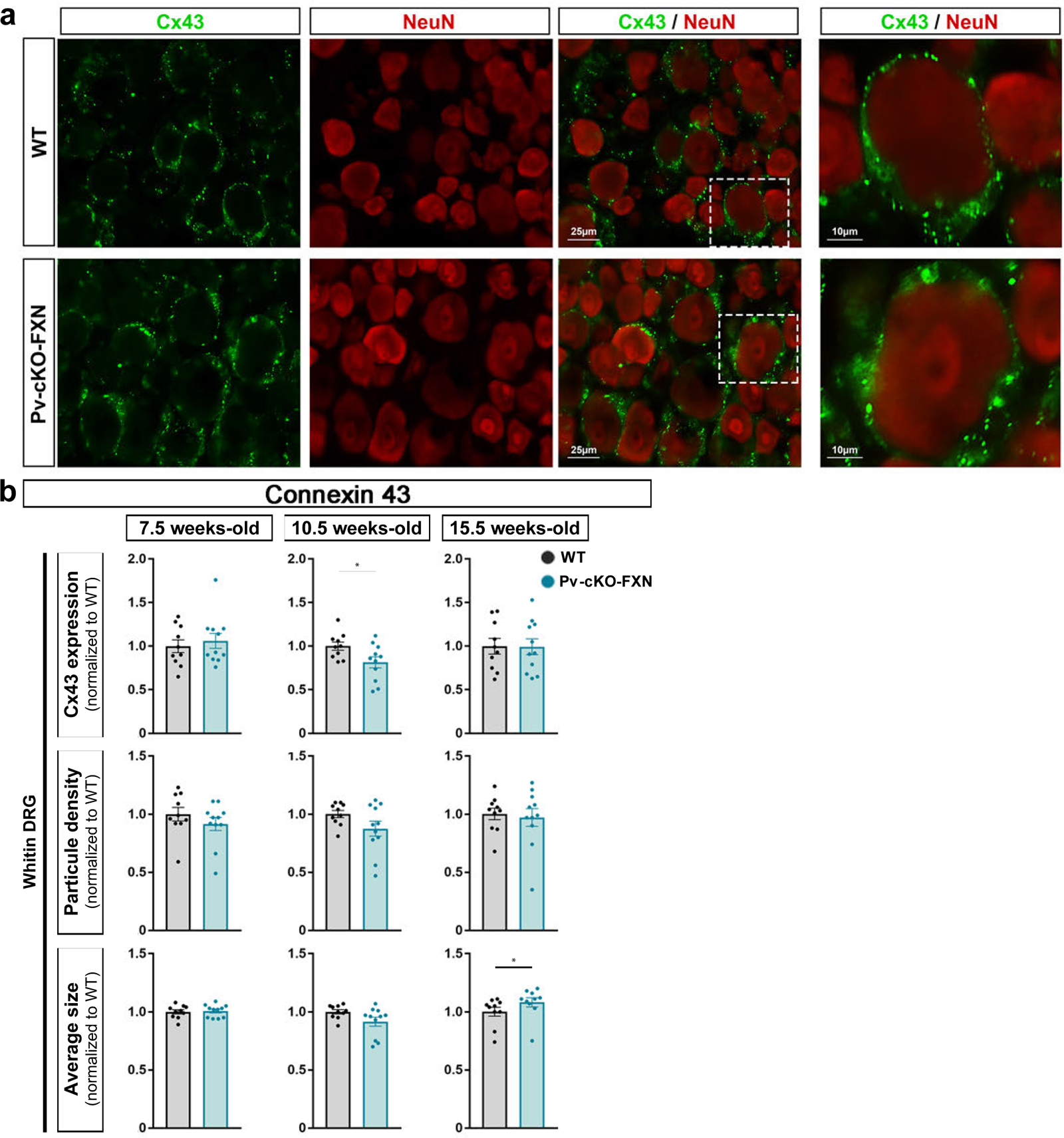
Immunohistochemistry of Cx43 and NeuN in DRG of control and Pv-cKO-FXN mice. (a) Representative images of immunofluorescence staining of DRG of WT and Pv-cKO-FXN animals at 10.5 weeks of age; with Cx43 (green) which labels SGC surrondding neurons marked with NeuN (red). Scale bar 25µm and 10µm for magnification. (b) Quantification mean density, number of particule and size measurements for Cx43-immunostaining across the entire DRG. All data shown symptomatic stages (7.5 10.5 15.5 week-old); are mean ± SEM. An unpaired parametric T-test has been performed. ***p<0.005; n=10 WT mice ; n=11 Pv-cKO-FXN mice.

Finally, to identify candidate molecular candidates and to better understand the molecular progression related to ganglia pathophysiology, we analyzed gene transcription at late 15.5 week-old experimental stage. At this stage, we observed 16 DEGs: the majority (13 genes) was upregulated, consistent with *Jun* upregulation, and only 3 genes were downregulated (***Figure 7***). Notably, and in agreement with the inflammatory program initiated and amplified in previous stages, several genes implicated in activation of immune responses were upregulated, including *Lpar6*, *Ccl12*, *Tlr7*, *Sphk1*, and *Csf1*. *Lpar6* gene product is a lysophosphatidic acid receptor promoting BBB permeability^79^ and is expressed in DRG endothelial cells^24^. *Ccl12* is an orthologous gene of *Ccl2* and encodes a potent monocyte chemoattractant protein mediating macrophage and T cells recruitment to sites of inflammation produced by tissue injury^80^. *Ccl12* is known to expressed in DRG macrophages^24^. *Tlr7* gene product (Toll Like Receptor 7; TLR7) is an innate immune receptor that triggers the production of proinflammatory cytokines and interferons in immune cells ^81,82^. In mouse DRGs *Tlr7* is expressed in macrophages and SGCs and not in sensory neurons ^24,81^. Together these findings suggest that at 15.5 week-old stage, BBB remains permeable, M1 polarization and production of proinflammatory mediators are both still ongoing, which may all result in building a toxic microenvironment for proprioceptors. Furthermore, it is worth mentioning that SGC *Sphk1* and neuronal *Csf1* also remained upregulated at this stage, and concomitantly SGC started to show sign of reactivity as evidenced by an increased expression of the SGC reactivity marker *Gfap*^24,83–85^ (***Figure 7***). A greater reactivity of SGCs was reinforced by the fact that the size of Cx43 expressing particles was increased by 8% within DRGs of Pv-cKO-FXN mice compared to controls (***Figure 10b***). Such increase suggests that SGCs are interconnected by larger sized gap junctions, rather than normally sized gap junctions at late 15.5 week-old symptomatic stages. This could reflect an adaptive response to FXN deficiency in proprioceptors and contribute to neuronal dysfunction. Indeed, increased Cx43 expression level was reported in FA Human patients’ DRGs ^9^. Furthermore, there is recent evidence for a role of injury-induced upregulation of Cx43-mediated gap junctions in DRG SGCs that contributes to sensory neuron hyperactivity and pain hypersensitivity^86^. All together, these observations are consistent with the idea that SGC-macrophages and SGC-neuron interactions play central detrimental role in ganglionopathy progression at 15.5 week-old stages, which is consistent with concomitant dramatic sensorimotor deficits (***Figure 1***). Interestingly, the clustering of CD68 expressing macrophages around proprioceptors was no longer observed at 15.5 week-old stage (***Figure 8***) that may indicate a resolution of the homeostatic inflammatory recruitment response.

### Selective FXN loss in proprioceptors leads to transcriptional changes in genes related to myelination, neuronal activity, neurodegeneration and neuronal disorders

In order to provide insight into how FXN deficient proprioceptors causes ganglionopathy initiation and progression, we next extracted relations between i) functions revealed through bioinformatic pathway analysis, ii) specific DEGs involved in neuronal activity or function, and previous literature, validating genes for certain cellular or functional phenotypes.

We first identified DEGs known to be involved in myelination, which may be relevant to FA since patients’ neuropathological examination shows depletion of large proprioceptive myelinated axons^87^. At the early behaviorally asymptomatic 3.5 week-old stage, we observed a downregulation of *Pmp22, Mbp, Bcas1, Utg8a* genes in DRGs of Pv-cKO-FXN compared to WT mice (***Figure 3***). These four genes are expressed in DRG SGCs and Schwann cells^24,29,85,88^ and have well established roles in myelin synthesis ^89–91^. Their downregulation suggests altered myelogenesis and/or defective myelin repair, which is consistent with the previous observation that at 3.5 weeks of age, sciatic nerves of Pv-cKO-FXN mice already displayed signs of myelin inner tongue abnormalities and beginning of neuropathy^14^. Furthermore, abnormal expression of *Pmp22* or *Utg8a* produces neurodegenerative phenotypes resembling Charcot-Marie-Tooth or associated to Alexander’s diseases^88,92^.

Noticeably, at the 7.5 week-old stage, when macrophages invade DRG of Pv-cKO-FXN mice, we observed an upregulation of *Adamts1* and *Il-18* genes encoding a metalloproteinase and proinflammatory cytokine, respectively (***Figure 5***). In DRG, *Adamts1* is mainly expressed in endothelial cells and pericytes, and to a lesser extent in SGCs, while *Il-18* is equally found in macrophages and SGCs^24^. Remarkably, *Adamts1* is upregulated in Alzheimer’s and Pick’s diseases as well as Down-syndrome, and regarded as a disease biomarker^93^. Il-18 is enhanced in Alzheimer’s and Parkinson’s diseases, and is believed to be unfavorable to neuronal survival^94,95^. Consistent with findings in FA drosophila models^19,49^ and cerebral ataxia patients^50^, we also observed an increases *Igf1* expression in DRG of Pv-cKO-FXN mice, possibly modulating DRG sensory neuron Ca^2+^ homeostasis and promoting neuronal apoptosis^24,96–99^. These findings are indicative of neuronal dysfunction, and in support to the concomitant down-regulation of *Gjd2* expression (***Figures 4*, *5***), a feature previously described in DRGs of amyotrophic lateral sclerosis (ALS) patients and ALS mouse model^100^ as well as pain model^101^. These results are accompanied by increased *Slc6a1* and *Slc12a5* expressions, both known to also be upregulated in pain and involved in sensory neuronal hyperexcitability^102,103^.

At the late 10.5 week-old stage, other upregulated genes in Pv-cKO-FXN mice, including *Lpl*, *Sphk1*, *Islr2* and *Csf1* were also previously associated with neurodegenerative diseases (***Figure 6***). In particular, *Lpl* is upregulated in Alzheimer’s disease^69^. *Sphk1* is overexpressed in neuroinflammatory disorders, including Alzheimer’s and Parkinson’s diseases^42,70^. *Islr2* neuronal gene is related to neuronal control of axon extension, and an increase of its expression is associated with neuropathic pain^104,105^. In addition, overexpression of *Csf1* participates in neuronal hyperexcitability in a pain context^72,106^.

At the 15.5 week-old stage, *Lpar6* was upregulated in DRG of Pv-cKO-FXN mice (***Figure 7***). Interestingly this gene was found overexpressed in the early presymptomatic and symptomatic stage of Huntington’s disease patients^107,108^. We also observed 3 downregulated genes (*Nefh*, *Cnr1*, *Cux2*) which are considered to be neuronal markers^109–111^. The neurofilament *Nefh* and *Cnr1* genes are known to be deregulated in Charco-Marie-Tooth’s disease and ALS as well as in pain, respectively^110,112^.

All together, these findings connect FA neurodegenerative disorder with molecular signaling pathways known to be involved in other neurodegenerative diseases (but other pain), supporting the growing evidence that different neurodegenerative diseases may overlap at multiple molecular and pathway levels ultimately leading to neuronal death.

## Discussion

Our study revealed that FXN deficiency selectively in proprioceptors causes major changes in inflammatory gene transcription as well as in macrophage and SGC gene and anatomical responses, highlighting molecular and cellular pathways that were sequentially altered, thus representing potential biomarkers and temporal signatures of FA ganglionopathy progression. More specifically, our analysis identified changes in expression of various genes related to distinct networks, pathways and biological/pathological functions, which for some of them are similar to those reported in FA patients or animal models, or other neurodegenerative diseases. We observed that FXN deficiency in proprioceptors is sufficient to induce i) gene modulations involved in general multicellular inflammatory processes (*Tspo*), immune cell infiltration, macrophage polarization, macrophage recruitment, apoptosis, fatty acid metabolism, myelination, neurodegeneration, and ii) macrophage and SGC anatomical and number changes within DRG and around proprioceptors. These abnormalities are also accompanied by transcriptional changes indicative of delayed/altered maturation of macrophages at asymptomatic ganglionopathy stage, and on the contrary, early elimination of the SGCs surrounding FXN deficient proprioceptors at late ganglionopathy stage. Given the well-established role for macrophages and inflammatory mediators^34,113^ and SGCs^114^ in regulating sensory neurons functions, as well as the similarity between human and mouse SGCs^27^, we believe that our observation of functional and anatomical disruptions of neuron-macrophages-SGC communications may have important contribution to proprioceptor dysfunctions and degeneration in FA ganglionopathy. Thus, our study will contribute to a better understanding of the mechanisms by which FXN deficiency leads to the different phenotypes observed in FA, which is a major goal of current research. As an example of unexpected but interesting discovery, our unbiased pathway analysis revealed evidence for impairments in fatty acid metabolism function. Although little is known about lipid homeostasis in FA, levels of phosphatidylethanolamine, phosphatidylserine and linoleic acid are decreased in patients’ plasma samples^115–117^ and lipid droplets are found in glial cells of a FA drosophila model ^118^. Moreover, lipid metabolism function has recently been shown to be important in the communication of SGCs to injured neurons^119^. Thus our results support an involvement of glial cells (*e.g*. DRG SGCs and/or Schwann cells) and lipid metabolism/peroxidation in the generation of FA ganglionopathy, an area of research we plan to investigate further using single cell RNASeq using our Pv-cKO-FXN mouse model in comparison with conventional FA models displaying ubiquitous FXN knockdown.

The use of the conditional Pv-cKO-FXN mouse model for modeling progressive ganglionopathy and sensory neuropathy represented a great advance, as it is the only tool available so far, that has allowed us to successfully test whether FXN deficiency in proprioceptors is causal to the inflammatory and SGC responses reported in FA patients’ DRGs. Here we first confirmed in our hands that Pv-cKO-FXN mice develop a progressive loss of coordination. We also completed behavioral characterization by measuring reflexes using the hindlimb extension reflex test. This simple test that does not require any animal training nor equipment has proven to be more sensitive to detect early behavioral deficits than both two-limb hanging and static bar tests. Therefore, investigators in the field may want to systematically add such broadly applicable test to batteries of behavioral tests routinely used for FA animal model evaluation.

Additionally, the selective deficiency of FXN in proprioceptors constitutes both the strength and the weakness of the Pv-cKO-FXN model. Indeed, this model cannot be used to study the impact of other cell FXN deficiency on ganglionopathy progression as well as on proprioceptor vulnerability and plasticity^14^. However, the fact that our findings show that selective FXN deficiency in proprioceptors is sufficient to recapitulate several key phenotypes observed in patients (macrophages clustering around unhealthy sensory neurons, increase in Cx43 expression, TSPO increase in nervous system^9,120^ as well as some gene transcript changes reported in ubiquitous FA models (*Tspo, Igf1 and Stab1*), suggests that FXN downregulation in proprioceptors is central to cause the main cellular and molecular phenotypes underlying DRG ganglionopathy initiation and progression. Our finding promise to have profound implications for FA disease understanding at the basic level but also for identifying or confirming signaling molecules and pathways to develop novel therapeutic strategies to cure or alleviate the disease.

Finally combining the use of this Pv-cKO-FXN model with the use of the nCounter technology and the *Glial Profiling panel*, covering 770 genes across 50 pathways has allowed for a comprehensive study of the complex interplay between neurons, peripheral immune cells, and SGCs. Our RNA profiling will be made fully available, which we hope will be of help to the field.

Among the DEGs, one of our major findings was the upregulation of *Tspo* from the behaviorally asymptomatic to late symptomatic stages in Pv-cKO-FXN mice, revealing *Tspo* (and its product) as an early biomarker of ganglionopathy. This is consistent with the facts that i) in DRGs, *Tspo* is widely expressed in basically all cell types^24^, ii) in FA patients radiolabeled ligands of TSPO^120^ and blood cell *Tspo* transcripts both show progressive increases over the course of the disease^18^, and iii) ubiquitous FA animal model also show *Tspo* upregulation^19^. Interestingly upregulation of TSPO and ADAMTS1 protein expressions were reported in other neurodegenerative diseases, including ALS, Huntington’s disease, Parkinson’s disease and Alzheimer’s disease^16,17,20,93^, highlighting both TSPO and ADAMTS1 as common biomarkers of several neurodegenerative diseases, including FA.

Furthermore, despite differences between FA models (mouse, drosophila), the approaches used for silencing/reducing *Fxn* expression (CRISPR-Cas9, Cre-lox, or shRNA), the tissues studied (DRG, brain, heart), our study identified several DEGs that are common between animal models and patients as well as between different animal models. For instance, in addition to *Tspo*, we observed that *Emp1*, *Igfbp5*, *Erg1*, *Itma2a* and *Smad3* genes are differentially expressed in Pv-cKO-FXN mouse DRGs as well as in patients’ blood cells^18,121^. Similarly, *Erg1, Itgam, C1qc, C3ar1, Clec7a, Cd68, Aif1, Gpnmb, Smad3* genes were differentially expressed in Pv-cKO-FXN mouse DRGs and other animal models^19,40,122^.

Finally, if we compare closely our results to available transcriptional analysis from another mouse FA mouse^40^, we did not find many common DEGs. The most rational, and not exclusive, explanations for this discrepancy are two folds: i) Pv-cKO-FXN model leads to FXN knockout selectively in proprioceptors from embryonic stages to adulthood while the inducible FRDAkd model leads to knockdown in all cell types (starting in the adulthood); ii) all cell types may not have the same vulnerability to extinction or drastic downregulation of FXN expression and their vulnerability may also vary depending on the developmental stages at which FXN is becoming deficient^49^. For instance, DRG proprioceptors are known to express 44-66% of total DRG FXN^14^, which may explain in part their vulnerability to FXN deficiency, and thus their early and specific involvement in DRG ganglionopathy.

Another main finding of this study is related to differences in the sequential immune response activation that we observed between Pv-cKO-FXN and WT mice throughout the different experimental stages. In Pv-cKO-FXN mice, behaviorally presymptomatic (3.5 week-old) and early symptomatic (5.5 week-old) stages were marked by a downregulation of many inflammatory related genes (*C3ar1, C1qc, Cd109, Chil1,Clec7a, Emp1, Fcgr1,Itm2a, Icam2, , Smad3, Spp1, Tlr2*), including a number of macrophage marker genes (*Aif1, Cd14, Cd68, Cx3cr1, Gpr34, Itgam*). The major resident macrophage populations in tissues are established before birth and are maintained thereafter into adulthood, independent of turnover by blood monocytes^123^. During embryonic development, CX3CR1 expressing macrophages invade tissues, including DRGs^124^. Although it is known that adult neurons can secrete CXCL1 (*i.e*. CX3CR1 ligand)^77,78^, it remains to be determined whether and how sensory neurons and macrophages interact during DRG development. Our findings are consistent with the existence of a proprioceptor-macrophage dialog (perhaps involving CX3CR1/CXCL1 pathway, and also SGCs the closest proprioceptors neighbors) that may occur during embryogenesis or early developmental stages in order to attract macrophages to become the DRG resident macrophages found in juvenile (*e.g.* 3.5 to 5.5 week-old stages) and adult stages (*e.g.* 7.5 to 15.5 week-old stages). One can speculate that FXN deficient proprioceptors are not healthy/mature enough to optimally produce and secrete chemoattractant mediators during embryogenesis.

In addition, at intermediate 7.5 week-old stage, there was a switch with a sharp upregulation in inflammatory genes, displaying mostly signature of M2 macrophage genes (*Igf1, Fcrls, Ms4a4*), which was accompanied by the anatomical recruitment of CD68 expression macrophages within whole DRGs. At the next symptomatic 10.5 week-old stage, we observed upregulation of other inflammatory genes (*Cd84, Csf1, Cx3cl1, Irf8, Lpl, Lrrc25, Mafb, Sphk1, Stab1*), some of them being known to have functions in efferocytosis (*i.e.* clearance of apoptotic cells by phagocytes for the maintenance of tissue homeostasis; *Stab1, Mafb*). This was concomitant with the increased *Cx3cl1* gene transcript, which product is known as a soluble “find me” factor released by neurons and apoptotic cells^77,78,125^, as well as the upregulation of macrophage marker genes (*Cd84, Lccr25, Mafb, Stab1*) involved in recognizing phophatidylserine to phagocyte apoptotic cells^63,65^. These transcriptomic results were further substantiated by the detection of CD68 expressing macrophages aggregated specifically around proprioceptors, which was paralleled with a decreased number of SGCs around proprioceptors. Given that i) at 10.5 and 15.5 week-old stages, we did not observe any obvious decrease in neuronal density (all neurons, proprioceptors or nociceptors); ii) at 10.5 week-old we found a decreased number of SGCs surrounding proprioceptors in Pv-cKO-FXN compared to WT mice; iii) at 15.5 week-old stage the number of SGCs in WT mice became as decreased as in Pv-cKO-FXN mice; together our finding suggest that at 10.5 week-old stage, macrophages surrounding FXN deficient proprioceptors may have phagocytosed some of their surrounding SGCs. One can speculate that there is a normal physiological SGC developmental program, that remains to be clearly determined, leading to removal of some supernumerary SGCs surrounding large sensory neurons and occurring at the young adulthood periods (∼10.5 weeks of age). In the case of FXN deficient unhealthy proprioceptors, interactions between proprioceptors and SGCs may be disrupted causing this SGC developmental program to be activated several weeks earlier (at 10.5 rather than 15.5 week-old). In the future it will be important to study further SGC (but also macrophage) development programs. This knowledge will be relevant to understand better physiological or abnormal molecular and cellular changes occurring in FA ganglionopathy or other neurological chronic DRG disorders. We ambition to test whether such programs take place using scRNASeq and DRGs from different development stages (from embryonic to late adulthood).

At late symptomatic 15.5 week-old stage, several genes implicated in activation of immune responses were upregulated in DRG of Pv-cKO-FXN. However, upregulation of two specific genes retained our attention as possible markers for a new inflammatory response shift. First, upregulation of the macrophage *Ccl12* gene (orthologous to *Ccl2*) encoding a chemoattractant cytokine very common in the recruitment of immune cells^126–128^. Second, upregulation of the specific SGC *Gfap* gene, suggesting that SGCs have entered into a reactivity state^24,83–85,119^. At this 15.5 week-old stage, CD68 expressing macrophages are no longer aggregated around FXN deficient proprioceptors, indicating that they indeed mediate different function (perhaps promoting global toxic proinflammatory inflammation throughout whole DRGs) compared to 10.5 week-old stage. Our findings suggest also that the remaining SGCs that envelop proprioceptors have lost important physiological functions and/or gained toxic functions related to their activated state. Such SGC reactivity is reminiscent of the enhanced GFAP expression level found in pain models^96,129,130^. Further studies carried out at later symptomatic stages^14^ (*e.g.* 20 week-old) may allow us to determine whether reactive SGCs will proliferate around FXN deficient proprioceptors^9^ (as found in patients’ DRGs of late FA stage).

Recent studies using FA ubiquitous mouse models have reported that endoplasmic reticulum (ER) Ca^2+^ content was pathologically reduced, mitochondrial Ca^2+^ uptake was impaired, and cytosolic Ca^2+^ concentration was enhanced in neurons^131–133^. This Ca^2+^ dyshomeostasis is due to the excess of oxidative stress under FA conditions and subsequent aberrant modulation of key players at the ER and mitochondrial levels that maintain physiological Ca^2+^ homeostasis. Interestingly at the late symptomatic 15.5 week-old, an upregulation of the neuronal *Ip3r1* transcript was detected. This gene product is located that the ER plasma membrane and is involved in the maintenance of mitochondrial and cytosolic Ca^2+^ concentrations. Together these observations suggest the possibility that IP3R1 is upregulated in FXN deficient proprioceptors of Pv-cKO-FXN mice, contributing to neuronal Ca^2+^ dyshomeostasis and thus to neuronal dysfunction. Consistent with this idea, neuronal Ca^2+^ dynamic alterations are known to be associated with other neurodegenerative diseases, including Alzheimer’s, Parkinson’s and Huntington’s diseases^134^. Thus, one of the next outstanding questions in the field is whether or not Ca^2+^ homeostasis and dynamics are altered in FXN deficient proprioceptors in living animal. This has not yet been tested while Ca^2+^ dyshomeostasis in proprioceptors represents an obvious and important candidate for inducing proprioceptor dysfunction and degeneration. Therapeutic strategies to find a treatment for FA are currently being developed to target proteins responsible for Ca^2+^ overload in DRG sensory neurons^133,135^.

In conclusion, our study provides, for the first time, evidence that FXN deficiency in proprioceptor is causal to the initiation and progression of both inflammatory and SGC responses observed in patients’ DRG ganglionopathy. In addition, it reveals molecular networks and pathways that are sequentially activated over disease progression and may represent potential biomarkers or signatures of the different disease stages. Finally, our results uncover important defects in neuron-macrophage-SGC anatomical association, suggesting that manipulation of both macrophages and SGC function, their revealed molecular signaling maybe worthy of consideration as a step towards strategies to restore/alleviate proprioceptive neuronal functions. Given the availability of a plethora of pharmacological tools targeting glia and macrophage-mediated inflammation, there is hope to succeed dampening inflammatory mediator-mediated aberrant activation of proprioceptor ionotropic or metabotropic receptors.

## Author contributions

PM and CA designed the experiments. PM performed most of experiments, analyzed and graphed the results. LW helped and advised about RNA purification. HP shared and advised about the Pv-cKO-FXN mouse model, and provided some DRG samples in the early stages of this project in order to obtain preliminary results before the mouse colony was fully amplified in CA’s laboratory. CA conceived, administrated and supervised the project. PM and CA wrote the manuscript. All authors read and commented the article.

## Acknowledgments

We gratefully acknowledge Sophie Guinoiseau for genotyping, perfusing and dissecting DRGs from the majority of mice used in the study; N. Diedhiou for perfusing and dissecting some DRGs used for preliminary immunohistochemistry experiments; Julia Weng and Trimitie Godreau for helping with SGC quantification; S. Lefebvre for advising with RNA purification; M. Rodero for useful discussions about genomic data analysis; Nicolas Cagnard from the Imagine Bioinformatic core facility for his great help in RNA profiling analysis; J.M. Andrieu for sharing laboratory spaces; F. Charbonnier for sharing pieces of equipment, and the BioMedTech mouse core facility for hosting our mouse colony.

---

**Figure S1.**
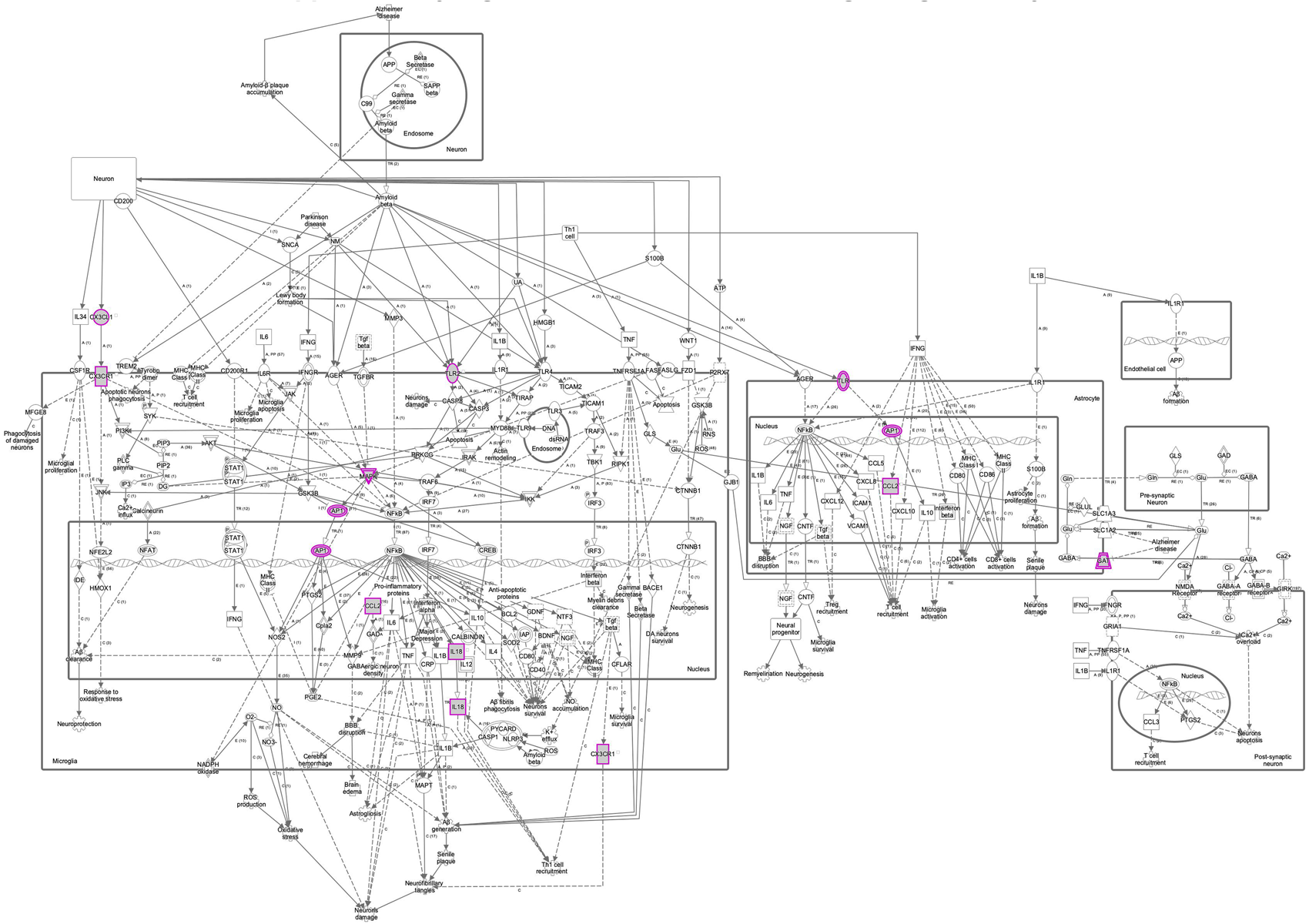
Neuroinflammation Signaling Pathway.

**table S1.** Antibody.

| Primary antibodies |  |  |  |
| --- | --- | --- | --- |
| Name | Company/Cat# | Species | Dilution |
| CD68 | Serotec/MCA1957 | Rat | 1:2'000 |
| GLAST | Frontier Institute/Af660 | Rabbit | 1:5'000 |
| Connexin 43 | Abcam/ab11370 | Rabbit | 1:1'000 |
| NeuN | Milipore/ABN90 | Guinea Pig | 1:1'000 |

| Secondary antibodies |  |  |  |
| --- | --- | --- | --- |
| Name | Company/Cat# | Species | Dilution |
| Alexa Fluor<sup>®</sup> Anti-Rat 546 | Invitrogen/A-11081 | Goat | 1:1'000 |
| Alexa Fluor<sup>®</sup> Anti-Rabbit 488 | Invitrogen/A-11034 | Goat | 1:1'000 |
| Alexa Fluor<sup>®</sup> Anti-Guinea pig 488 | Invitrogen/A-11074 | Goat | 1:1'000 |

**table S2.** Extraction ARN.

| RNA Extraction |  |  |  |  |  |
| --- | --- | --- | --- | --- | --- |
| Stages | Quantity (ng/μL) | A260/280 | A260/230 | DV200 % | RIN |
| 15,5 weeks-old | 73 | 1,72 | 1,67 | 60 | 7,3 |
|  | 76 | 1,93 | 1,62 | 70 | 7 |
|  | 73 | 1,18 | 1,86 | 72 | 7,2 |
|  | 84 | 1,8 | 2,13 | 76 | 7,8 |
|  | 51 | 1,87 | 1,7 | 74 | 7,7 |
|  | 42 | 1,78 | 1,87 | 71 | 7,6 |
|  | 44 | 1,82 | 1,77 | 66 | 7,4 |
|  | 51 | 1,8 | 1,87 | 68 | 7,2 |
| 10,5 weeks-old | 136 | 1,73 | 1,79 | 70 | 7,3 |
|  | 44 | 1,88 | 2,01 | 75 | 7,1 |
|  | 76 | 1,85 | 2,06 | 74 | 7,2 |
|  | 101 | 1,82 | 2,11 | 76 | 7 |
|  | 61 | 1,84 | 2,4 | 80 | 7,3 |
|  | 50 | 1,88 | 1,81 | 78 | 7,6 |
|  | 64 | 1,88 | 2,05 | 81 | 7,8 |
|  | 32 | 1,85 | 1,97 | 70 | 7,3 |
| 7,5 weeks-old | 124 | 1,76 | 1,76 | 81,3 | 7,5 |
|  | 127 | 1,88 | 2,09 | 85 | 7,3 |
|  | 107 | 1,85 | 2,01 | 85 | 7,6 |
|  | 92 | 1,81 | 2,09 | 84 | 7,3 |
|  | 94 | 1,88 | 2 | 83 | 7,6 |
|  | 82 | 1,85 | 2,11 | 77 | 7,3 |
|  | 51 | 1,84 | 1,8 | 81 | 7,7 |
|  | 58 | 1,81 | 1,76 | 82 | 7,9 |
| 5,5 weeks-old | 76 | 1,86 | 1,94 | 79 | 7,9 |
|  | 37 | 1,8 | 1,62 | 75 | 7,4 |
|  | 56 | 1,89 | 1,88 | 73 | 7,7 |
|  | 41 | 1,84 | 1,77 | 77 | 7,7 |
|  | 27 | 1,77 | 1,8 | 75 | 7 |
|  | 120 | 1,97 | 2,155 | 84 | 7,4 |
|  | 55 | 1,83 | 1,64 | 80 | 7,5 |
|  | 53 | 2 | 1,75 | 71 | 7,9 |
| 3,5 weeks-old | 204 | 1,77 | 1,94 | 82 | 8,1 |
|  | 50 | 1,8 | 1,95 | 82 | 8,3 |
|  | 167 | 1,77 | 1,85 | 78 | 8 |
|  | 40 | 1,85 | 1,74 | 92 | 8 |
|  | 37 | 1,87 | 1,7 | 91 | 7,8 |
|  | 42 | 1,8 | 1,7 | 91 | 7,8 |
|  | 59 | 1,77 | 2,13 | 90 | 7,8 |
|  | 114 | 1,88 | 1,85 | 83 | 7,7 |

**table S3.** BEHAVIOR.

| Hindlimb extension reflex |  |  |  |  |  |  |  |  |  |
| --- | --- | --- | --- | --- | --- | --- | --- | --- | --- |
| Number of mice |  | 2 way ANOVA |  |  |  | Bonferroni post-tests |  |  |  |
| WT | 10 | Source of Variation | % of total variation | P value |  | WT vs Pv-cKO-FXN |  |  |  |
| Pv-cKO-FXN | 11 | Interaction | 21 | P <0,0001 | **** | Stages | t | P value |  |
|  |  | Stage | 22,97 | P <0,0001 | **** | 3.5 | 0 | P >0,9999 | ns |
|  |  | Genotype | 31,35 | P <0,0001 | **** | 5.5 | 1,818 | 0,361 | ns |
|  |  |  |  |  |  | 7.5 | 4,746 | P <0,0001 | **** |
|  |  |  |  |  |  | 10.5 | 10,4 | P <0,0001 | **** |
|  |  |  |  |  |  | 15.5 | 10,93 | P <0,0001 | **** |
| Twolimb hanging |  |  |  |  |  |  |  |  |  |
| Number of mice |  | 2 way ANOVA - Falling |  |  |  | Bonferroni post-tests - Falling |  |  |  |
| WT | 10 | Source of Variation | % of total variation | P value |  | WT vs Pv-cKO-FXN |  |  |  |
| Pv-cKO-FXN | 11 | Interaction | 27,87 | P <0,0001 | **** | Stages | t | P value |  |
|  |  | Stage | 31,25 | P <0,0001 | **** | 5.5 | 0,1238 | >0,9999 | ns |
|  |  | Genotype | 30,38 | P <0,0001 | **** | 7.5 | 1,317 | 0,7661 | ns |
|  |  |  |  |  |  | 10.5 | 8,96 | P <0,0001 | **** |
|  |  |  |  |  |  | 15.5 | 14,24 | P <0,0001 | **** |
|  |  | 2 way ANOVA - Grasping |  |  |  | Bonferroni post-tests - Grasping |  |  |  |
|  |  | Source of Variation | % of total variation | P value |  | WT vs Pv-cKO-FXN |  |  |  |
|  |  | Interaction | 11,58 | 0,0003 | *** | Stages | t | P value |  |
|  |  | Stage | 32,85 | P <0,0001 | **** | 5.5 | 0,01018 | >0,9999 | ns |
|  |  | Genotype | 15,7 | P <0,0001 | **** | 7.5 | 0,8814 | >0,9999 | ns |
|  |  |  |  |  |  | 10.5 | 4,356 | 0,0002 | *** |
|  |  |  |  |  |  | 15.5 | 5,359 | P <0,0001 | **** |
|  |  | 2 way ANOVA - Crossing |  |  |  | Bonferroni post-tests - Crossing |  |  |  |
|  |  | Source of Variation | % of total variation | P value |  | WT vs Pv-cKO-FXN |  |  |  |
|  |  | Interaction | 4,143 | 0,1071 | ns | Stages | t | P value |  |
|  |  | Stage | 31,83 | P <0,0001 | **** | 5.5 | 0,9326 | >0,9999 | ns |
|  |  | Genotype | 13,8 | P <0,0001 | **** | 7.5 | 1,183 | 0,9612 | ns |
|  |  |  |  |  |  | 10.5 | 3,471 | 0,0034 | ** |
|  |  |  |  |  |  | 15.5 | 3,489 | 0,0032 | ** |
| Static bar |  |  |  |  |  |  |  |  |  |
| Number of mice |  | 2 way ANOVA - Falling |  |  |  | Bonferroni post-tests - Falling |  |  |  |
| WT | 10 | Source of Variation | % of total variation | P value |  | WT vs Pv-cKO-FXN |  |  |  |
| Pv-cKO-FXN | 11 | Interaction | 37,59 | P <0,0001 | **** | Stages | t | P value |  |
|  |  | Stage | 36,26 | P <0,0001 | **** | 5.5 | 0,2822 | >0,9999 | ns |
|  |  | Genotype | 23,03 | P <0,0001 | **** | 7.5 | 1,009 | >0,9999 | ns |
|  |  |  |  |  |  | 10.5 | 4,291 | 0,0002 | *** |
|  |  |  |  |  |  | 15.5 | 21,64 | P <0,0001 | **** |
|  |  | 2 way ANOVA - Distance |  |  |  | Bonferroni post-tests - Distance |  |  |  |
|  |  | Source of Variation | % of total variation | P value |  | WT vs Pv-cKO-FXN |  |  |  |
|  |  | Interaction | 32,02 | P <0,0001 | **** | Stages | t | P value |  |
|  |  | Stage | 31,38 | P <0,0001 | **** | 5.5 | 0,07445 | >0,9999 | ns |
|  |  | Genotype | 24,21 | P <0,0001 | **** | 7.5 | 0,3802 | >0,9999 | ns |
|  |  |  |  |  |  | 10.5 | 5,544 | P <0,0001 | **** |
|  |  |  |  |  |  | 15.5 | 14,44 | P <0,0001 | **** |
|  |  | 2 way ANOVA - Crossing |  |  |  | Bonferroni post-tests - Crossing |  |  |  |
|  |  | Source of Variation | % of total variation | P value |  | WT vs Pv-cKO-FXN |  |  |  |
|  |  | Interaction | 28,31 | P <0,0001 | **** | Stages | t | P value |  |
|  |  | Stage | 27,84 | P <0,0001 | **** | 5.5 | 0,4565 | >0,9999 | ns |
|  |  | Genotype | 22,3 | P <0,0001 | **** | 7.5 | 0,01388 | >0,9999 | ns |
|  |  |  |  |  |  | 10.5 | 5,985 | P <0,0001 | **** |
|  |  |  |  |  |  | 15.5 | 10,54 | P <0,0001 | **** |

**table S4.**
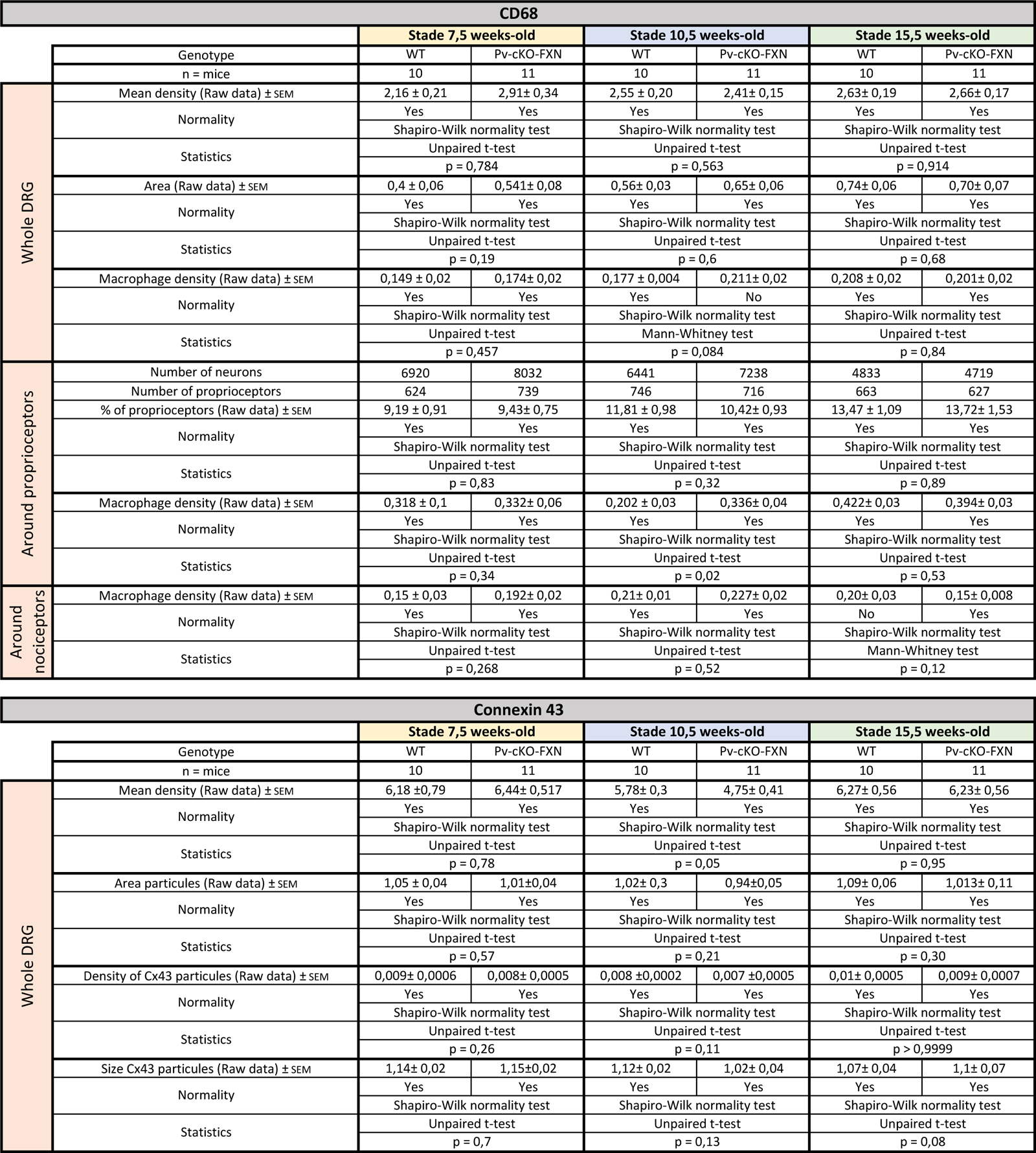
Immunohistochemistry #1.

**table S5.**
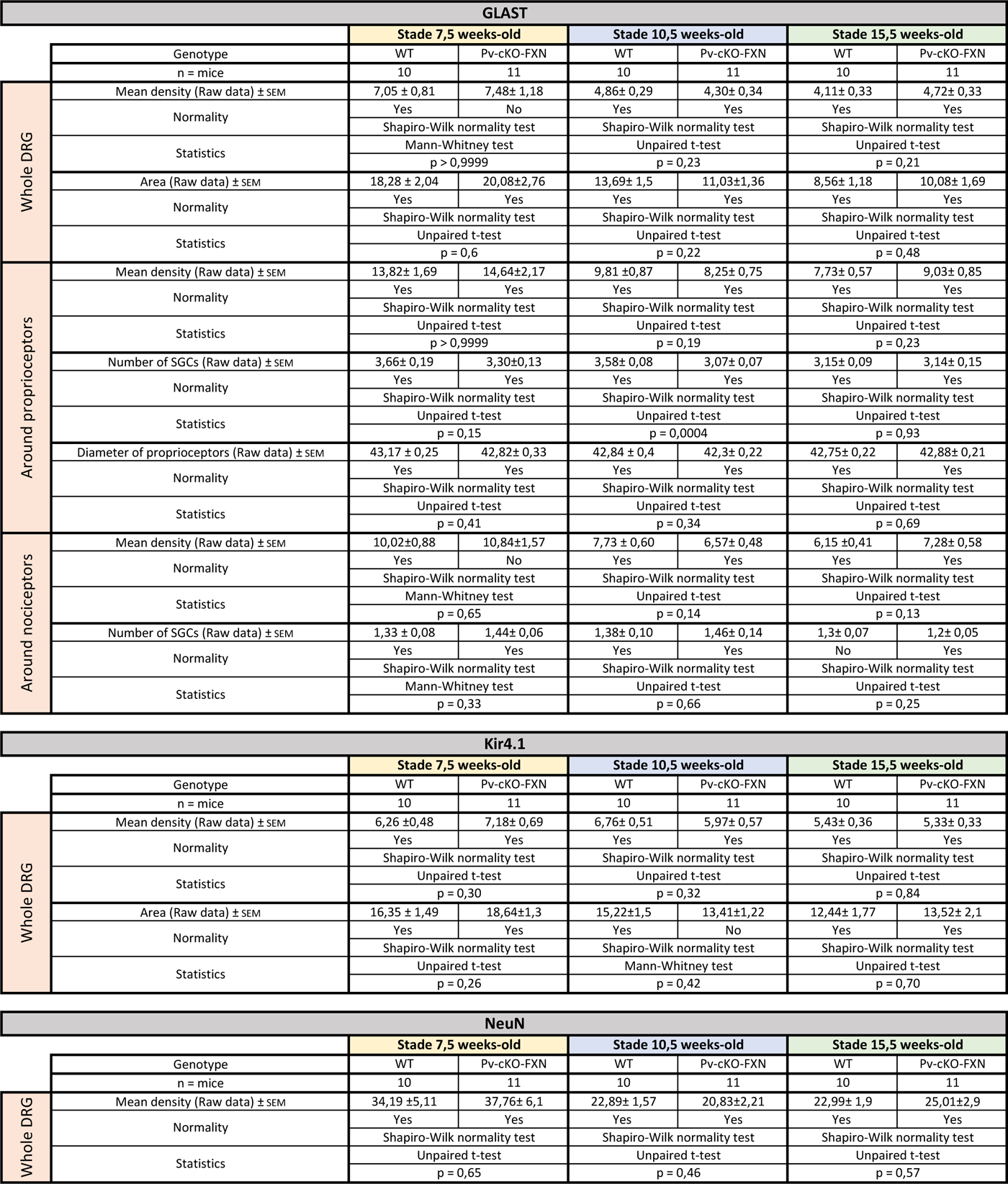
Immunohistochemistry #2.

